# Protein-primed DNA homopolymer synthesis by an antiviral reverse transcriptase

**DOI:** 10.1101/2025.03.24.645077

**Authors:** Stephen Tang, Rimantė Žedaveinytė, Nathaniel Burman, Shishir Pandey, Josephine L. Ramirez, Louie M. Kulber, Tanner Wiegand, Royce A. Wilkinson, Yanzhe Ma, Dennis J. Zhang, George D. Lampe, Mirela Berisa, Marko Jovanovic, Blake Wiedenheft, Samuel H. Sternberg

## Abstract

Bacteria defend themselves from viral predation using diverse immune systems, many of which sense and target foreign DNA for degradation^1^. Defense-associated reverse transcriptase (DRT) systems provide an intriguing counterpoint to this strategy by leveraging DNA synthesis instead^2,3^. We and others recently showed that DRT2 systems use an RNA template to assemble a *de novo* gene, leading to expression of an antiviral effector protein, Neo^4,5^. It remains unknown whether similar mechanisms of defense are employed by other DRT families. Focusing on DRT9, here we uncover an unprecedented mechanism of DNA homopolymer synthesis, in which viral infection triggers polydeoxyadenylate (poly-dA) accumulation in the cell to drive abortive infection and population-level immunity. Cryo-EM structures reveal how a conserved noncoding RNA serves as both a structural scaffold and reverse transcription template to direct hexameric complex assembly and RNA-templated poly-dA synthesis. Remarkably, biochemical and functional experiments identify conserved tyrosine residues within the reverse transcriptase itself that prime DNA synthesis, leading to the formation of high-molecular weight protein-DNA covalent adducts. Synthesis of poly-dA *in vivo* is regulated by the competing activities of phage-encoded triggers and host-encoded silencers of DRT9. Collectively, our work unveils a novel nucleic acid-driven defense system that expands the paradigm of bacterial immunity and broadens the known functions of reverse transcriptases.

## INTRODUCTION

Bacteria encode diverse mechanisms for antiviral defense, the majority of which degrade invading nucleic acids using either programmable, sequence-specific, or non-specific nucleases^1^. Also common among phage defense systems are enzymes that act on host nucleic acids — synthesizing second messenger molecules^6^, converting nucleotides into suicide/terminator analogs^7^, or depleting nucleotides altogether^8,9^. Against this backdrop, reverse transcriptase (RT) enzymes, which polymerize rather than degrade DNA, would seem unlikely contributors to antiphage defense. And yet, multiple families of RTs have recently been implicated in defensive functions spanning the adaptive and innate arms of the bacterial immune system^2–5,10,11^.

RTs associated with CRISPR-Cas comprise one family of defense-related systems, leveraging RNA-templated DNA synthesis to facilitate spacer acquisition directly from viral RNA transcripts^10^. But whereas CRISPR-associated RTs reverse transcribe foreign RNA, other RT systems encode and reverse transcribe their own non-coding RNA (ncRNA). Retron-encoded ncRNAs template the synthesis of hybrid RNA–DNA molecules — known as multicopy single-stranded DNA (msDNA) — that serve as antitoxins against host-encoded toxins^12^. Upon viral infection, a cascade of events leads to msDNA modification or degradation and toxin release, killing the infected cell and thereby preempting further viral propagation^12,13^. This strategy, known as abortive infection, protects the larger bacterial population at the cost of the infected cell, and is ubiquitous among innate immune mechanisms^1^. Abortive infection also characterizes a third group of antiviral RTs typified by AbiK, AbiA, and AbiP2, which, unlike CRISPR-associated RTs and retrons, have been shown to synthesize random DNA polymers independently of any RNA template^14,15^. Intriguingly, recent structural and biochemical studies revealed that DNA polymerization by Abi RTs is primed by a tyrosine residue within the RT itself and requires the formation of a homo-oligomeric RT complex^15,16^. These studies have demonstrated the presence of highly unusual biochemical behaviors across the Abi RT family, but how these activities relate to phage infection, in terms of their regulation and their effects on the phage and/or the host, remains unresolved.

The Abi RTs belong to a large and diverse family of systems collectively designated as the Unknown Group (UG), whose enigmatic functions highlight the underappreciated roles of RT-mediated defense in bacteria^17^. Pioneering work by Gao et al. and Mestre et al. revealed an expanded set of UG systems that function in antiviral immunity, termed defense-associated RTs (DRT)^2,3^. These studies identified a variety of DRT-associated enzymatic domains that have also been implicated in defensive roles in other immune pathways, but intriguingly, a subset of DRTs is not associated with any other protein domains^3^. We and others recently resolved the molecular mechanism of one such system, termed DRT2^4,5^. Like retron systems, DRT2 operons encode a conserved, structured ncRNA, which provides the template for complementary DNA (cDNA) synthesis by the RT. Remarkably, rolling circle reverse transcription of the ncRNA leads to the synthesis of concatemeric cDNA (ccDNA) products, assembling intact promoters and a nearly endless open reading frame (*neo*) spanning ccDNA repeats. During phage infection, ccDNA is transcribed and translated to yield Neo proteins that drive cell growth arrest in a mechanism akin to abortive infection. Altogether, these studies have provided an early glimpse into the unprecedented and unique molecular functions of UG-family RTs, while leaving unresolved whether similar mechanisms exist across the vast diversity of unexplored UG systems.

We set out to uncover conserved mechanisms of rolling circle reverse transcription that would yield RNA-templated *de novo* gene assembly in other bacterial immune systems. In this regard, we focused on DRT9, because of its nearby placement on the phylogenetic tree and its similar genetic architecture to DRT2 (**Fig. 1a**). What we instead uncovered was an altogether distinct mechanism of RT-mediated innate immunity. Here we show that DRT9 systems are triggered by phage infection to produce long, uninterrupted chains of polydeoxyadenylate (poly-dA). Poly-dA synthesis is primed by tyrosine residues within the RT itself, and is templated by a structured ncRNA containing a universally conserved poly-uridine tract. Cryo-electron microscopy (cryo-EM) data reveal extensive interactions between protein and RNA components that mediate the assembly of a hexameric RT-ncRNA complex, in which both the priming tyrosines and templating uridines reside proximal to the RT active site. The production of poly-dA is sensitively regulated by a ‘tug-of-war’ between host- and phage-encoded factors, and its accumulation in cells drives a growth arrest response consistent with abortive infection. In addition to revealing an elegant and intricate mechanism by which DNA synthesis confers antiviral immunity, our work highlights an unprecedented synthetic pathway and biological function of a nucleic acid homopolymer.

**Figure 1.**
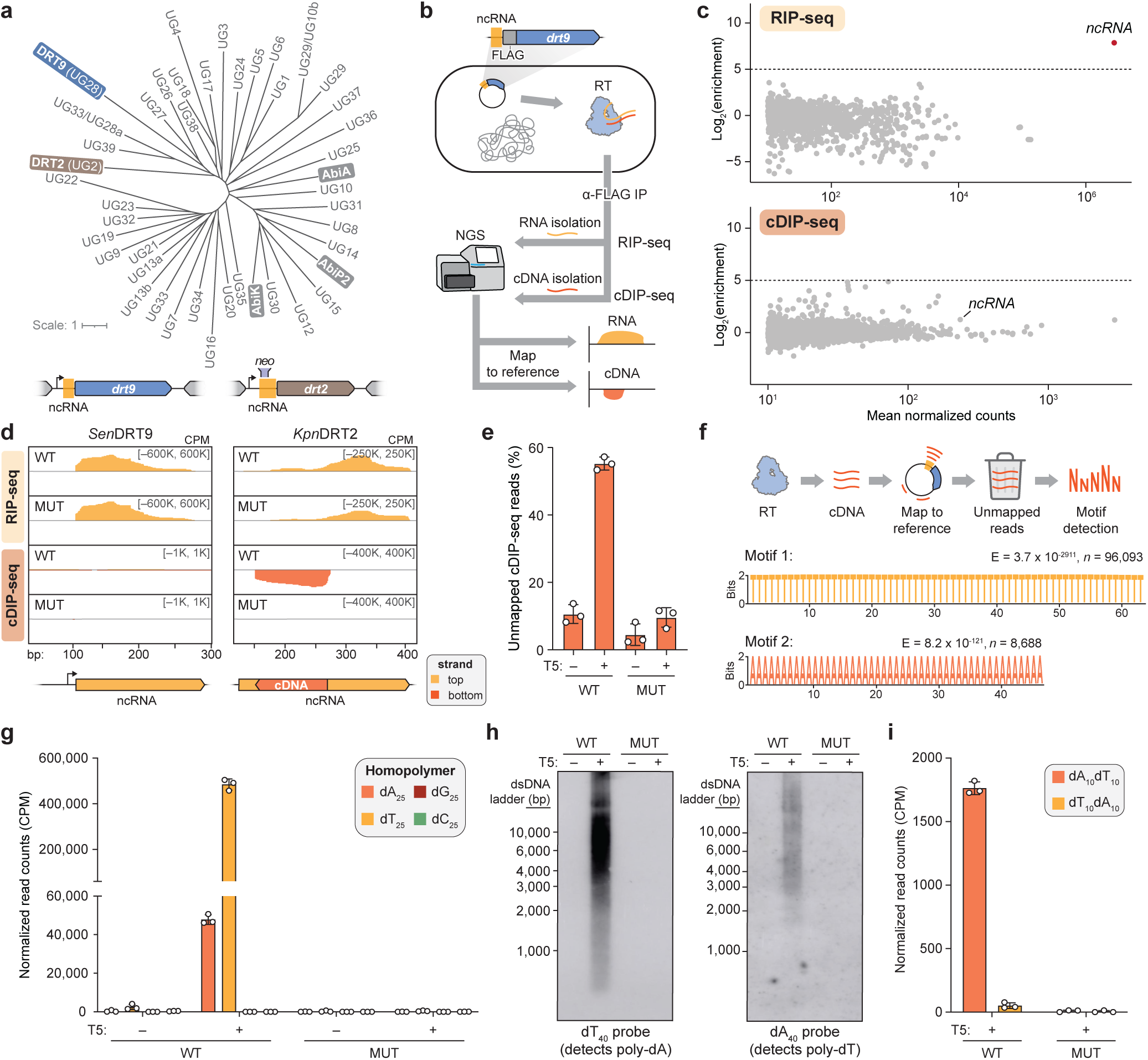
Systematic discovery of DRT9 reverse transcription substrates and products *in vivo*. **a,** Phylogenetic tree of bacterial reverse transcriptase (RT) homologs within the so-called Unknown Group (UG), which are broadly implicated in antiphage defense. DRT9 (UG28) and DRT2 (UG2) are highlighted, as are other systems that have been the subject of experimental studies; genetic architectures of archetypal DRT9 and DRT2 systems are shown below. The tree was adapted from Mestre *et al.*^3^ by collapsing each UG/DRT clade into a single representative. **b,** Schematic of RNA immunoprecipitation (RIP) and cDNA immunoprecipitation (cDIP) sequencing approaches to identify nucleic acid templates and products of FLAG-tagged reverse transcriptase (RT) from *Sen*DRT9. The plasmid-encoded immune system is schematized at the top. **c,** MA plots showing the RT-mediated enrichment of RNA (top) and DNA (bottom) loci from RIP-seq and cDIP-seq experiments, respectively, relative to input controls. Each dot indicates a transcript, and red dots indicate transcripts with log_2_(enrichment) > 5 and false discovery rate (FDR) < 0.05. **d,** RIP-seq and cDIP-seq coverage tracks (top to bottom) for either WT RT or a catalytically inactive RT mutant (MUT), in the presence of T5 phage infection, for *Sen*DRT9 (left) and *Kpn*DRT2 (right). Genomic locus schematics are shown below each graph, and data are normalized for sequencing depth and plotted as counts per million reads (CPM). Coordinates for DRT9 are numbered from the beginning of the *S. enterica*-derived sequence on the expression plasmid. **e,** Bar graph analyzing the percentage of unmapped cDIP-seq reads from WT or MUT *Sen*DRT9 cDIP-seq datasets in the absence or presence of T5 phage infection. Reads were mapped to a concatenated reference genome comprising the *E. coli* chromosome, phage genome, and expression plasmid. **f,** Schematic of unmapped read analytical pipeline (top), and MEME results that revealed poly-dT and poly-dA motifs enriched in unmapped reads from the WT + T5 cDIP-seq dataset in **e** (bottom). *E*, E-value significance of the motif; *n*, number of sites contributing to the motif. **g,** Bar graph of normalized dA_25_, dT_25_, dG_25_, and dC_25_ counts from WT and MUT cDIP-seq datasets in the absence or presence of T5 phage infection. **h,** Southern blot analysis of total DNA isolated from cells expressing WT or MUT *Sen*DRT9 in the absence or presence of T5 phage infection. Duplicate blots were probed with either oligo-dT_40_ (left) or oligo-dA_40_ (right) to detect poly-dA and poly-dT species, respectively. Sizes from a double-stranded DNA (dsDNA) ladder are marked, from the methylene blue-stained membrane after transfer (**Extended Data Fig. 2d**). **i,** Bar graph of chimeric dA_10_dT_10_ and dT_10_dA_10_ counts from WT and MUT cDIP-seq datasets in the presence of T5 phage infection. Data in **e,g,i** are shown as mean ± s.d. for n = 3 independent biological replicates.

## RESULTS

### Phage infection elicits synthesis of DNA homopolymers by DRT9

DRT9 systems are widely distributed across diverse bacterial taxa and exhibit a minimal genetic architecture consisting of a reverse transcriptase (RT) gene and an upstream ncRNA, occasionally in association with a SLATT domain-containing protein^3^ (**Extended Data Fig. 1a, Supplementary Table 1**). We selected 10 diverse systems for gene synthesis, expressed them in *E. coli*, and assessed their antiviral activity against a panel of well-characterized *E. coli* phages (**Extended Data Fig. 1b, Supplementary Table 2**). DRT9 systems from *Salmonella enterica* and *Photobacterium sanctipauli* (hereafter *Sen*DRT9 and *Psa*DRT9) robustly defended against multiple phages, particularly those from the *Tequatrovirus* and *Tequintavirus* genera, and this protection was abolished by mutations in conserved aspartate residues within the YADD active site (**Extended Data Fig. 1c,d**). Interestingly, epitope tagging at the C-terminus, but not the N-terminus, severely impaired *Sen*DRT9-mediated defense (**Extended Data Fig. 1e**). This observation was consistent with AlphaFold3 modelling of the *Sen*DRT9-encoded RT (*Sen*RT), which positioned the C-terminus proximal to the predicted catalytic residues, suggesting a potential role for the C-terminus in *Sen*RT enzymatic function (**Extended Data Fig. 1f**). Next, we tested the reverse transcriptase activity of N-terminally FLAG-tagged *Sen*RT, which retained wild-type (WT) defense function, by performing RNA immunoprecipitation (RIP) and cDNA immunoprecipitation (cDIP) sequencing^4^ in uninfected and T5 phage-infected cells (**Fig. 1b**). As seen with DRT2, the ncRNA encoded immediately upstream of the RT was strongly enriched by RIP-seq, however in contrast to DRT2, cDIP-seq analysis failed to detect evidence of reverse transcription upon phage infection (**Fig. 1c,d**).

Despite genetic evidence that reverse transcription is essential for DRT9-mediated defense, conventional analyses were unable to reveal the DRT9 cDNA product, leading us to wonder how we might uncover the bona fide reaction products. Inspired by reports that AbiK-encoded RTs synthesize untemplated, random DNA polymers^14,16^, we hypothesized that analyzing cDIP-seq reads that failed to map to the reference genome might reveal unexpected cDNA products. Inspection of alignment statistics revealed a striking increase in unmapped reads specifically in phage-infected cells expressing WT, but not mutant, *Sen*RT (**Fig. 1e**). This observation prompted us to manually inspect the unmapped reads, which uncovered an unanticipated result: long homopolymeric stretches of poly-deoxythymidylate (poly-dT) and poly-deoxyadenylate (poly-dA), a pattern further corroborated by unbiased motif detection analysis (**Fig. 1f, Extended Data Fig. 2a**). RIP-seq and cDIP-seq experiments with the related *Psa*DRT9 system similarly revealed strong enrichment of the upstream ncRNA, an absence of cDNA species mapping to the ncRNA, and a significant overrepresentation of poly-dT motifs within unmapped reads (**Extended Data Fig. 2b,c**). These results provide the first evidence of a reverse transcriptase naturally synthesizing homopolymeric DNA and implicate poly-dT and/or poly-dA products in the mechanism of antiphage defense.

We next developed an analytical approach to quantify reads containing homopolymers (≥25 nucleotides in length) across all cDIP-seq datasets for each of the four deoxynucleotides. Poly-dT and poly-dA were detected exclusively in phage-infected WT *Sen*DRT9 cells, with poly-dT outnumbering poly-dA by a factor of ∼10, while poly-dG and poly-dC products were not detectable above background (**Fig. 1g**). Nevertheless, given the potential for next-generation sequencing library preparation and/or sequencing artifacts to bias homopolymer identification and quantification, we turned to a classical molecular biology approach to assess poly-dT and poly-dA production in cells by an orthogonal method. Southern blotting experiments corroborated our cDIP-seq results, revealing abundant poly-dT and poly-dA species that were again restricted to phage-infected cells expressing WT *Sen*DRT9 (**Fig. 1h, Extended Data Fig. 2d**). Two additional observations from Southern blotting were notable. First, biotinylated oligo-dT and oligo-dA probes hybridized to cellular DNA species ranging from ∼1,000 to >10,000 bases in length, relative to a double-stranded DNA (dsDNA) ladder (**Fig. 1h**), indicating that *Sen*DRT9 synthesizes long DNA homopolymers. Second, poly-dA products were substantially more abundant than poly-dT products by Southern blotting, in stark contrast to the relative proportions observed by next-generation sequencing (**Fig. 1g,h**).

To further investigate the poly-dA vs. poly-dT discrepancy, we performed control experiments in which synthetic dT_100_ and dA_100_ oligonucleotides — mimicking DRT9 reaction products — were spiked into *E. coli* cell lysates at varying ratios prior to our standard DNA sequencing library preparation workflow (**Extended Data Fig. 2e**). These experiments revealed a strong bias against poly-dA capture and sequencing relative to poly-dT (**Extended Data Fig. 2f**), suggesting that Southern blot data more accurately reflect the true intracellular distribution of homopolymeric products during phage infection. High-throughput sequencing nevertheless provided insights that would have been challenging to detect otherwise, including the reproducible presence of chimeric species harboring dA_10_–dT_10_ transition sequences, albeit at significantly lower levels than poly-dA and poly-dT (**Fig. 1i**). Interestingly, these species were more abundant than their inverse counterparts (dT_10_–dA_10_), suggesting that initial cDNA synthesis by DRT9 may produce exclusively poly-dA, which occasionally serves as a template and a primer to generate a complementary poly-dT strand.

Collectively, these findings indicate that phage infection triggers the synthesis of poly-dA in the DRT9 immune response, while poly-dT likely arises from low-level second-strand synthesis, either by the RT itself or a host-encoded DNA polymerase. Of note, RNA-seq datasets failed to reveal any evidence of poly-A or poly-U species (**Extended Data Fig. 2g**), thus arguing against any coding role for DRT9 cDNA synthesis — a notable divergence from DRT2. This observation, along with the fundamentally distinct nature of the cDNA sequence, underscores the functional and molecular diversity of DRT enzymes and their reverse transcription products.

### Non-coding RNA sequence determinants of DNA homopolymer synthesis

How can a reverse transcriptase synthesize kilobase-length polymers composed almost exclusively of adenosine nucleotides? We initially considered the hypothesis that the RT might directly select dATP substrates through protein-mediated recognition while remaining otherwise template-independent, akin to the reported mechanism of untemplated DNA synthesis by AbiK^16^. However, closer inspection of the ncRNA sequence and predicted secondary structure — as informed by evolutionary covariation and conservation across 201 diverse DRT9 ncRNA sequences — suggested an alternative hypothesis. In addition to conserved stem-loop (SL) features that loosely resemble the overall architecture of DRT2-associated ncRNAs, the DRT9 ncRNA exhibits a universally conserved stretch of uridine residues positioned similarly to the reverse transcription start site of the DRT2 ncRNA (**Fig. 2a, Extended Data Fig. 3a,b**). We therefore envisioned that stable positioning of this uridine-rich template region within the RT active site could drive highly processive synthesis of complementary poly-dA chains, potentially via iterative realignment of the 3′ end of the growing poly-dA strand at invariant uridine nucleotides.

**Figure 2.**
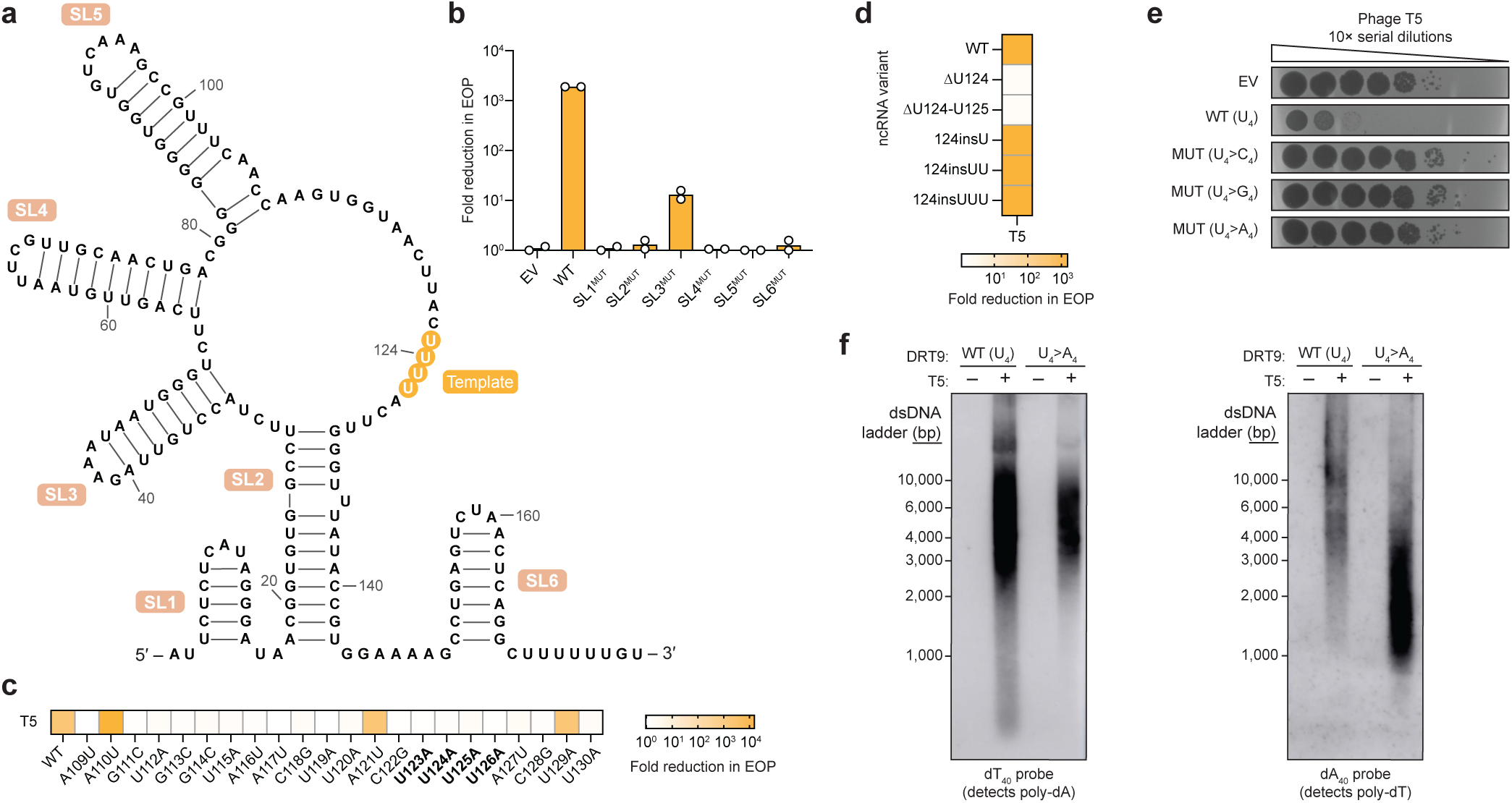
ncRNA sequence determinants of *Sen*DRT9-mediated phage defense and poly-dA synthesis. **a,** Predicted secondary structure of the *Sen*DRT9 ncRNA. Stem-loop (SL) regions and uridine nucleotides implicated in RNA-templated DNA synthesis are labeled; coordinates are numbered based on the mature ncRNA species identified by RIP-seq. **b,** Bar graph quantifying *Sen*DRT9 defense activity against T5 phage for scrambled ncRNA SL mutants (MUT), quantified as the fold reduction in efficiency of plating (EOP) relative to an empty vector (EV) control. Data are from n = 2 technical replicates. **c,** Heat map quantifying *Sen*DRT9 defense activity for the indicated ncRNA point mutations, quantified as the fold reduction in EOP relative to an empty vector (EV) control. Uridine nucleotides implicated in RNA-templated DNA synthesis are highlighted in bold; data are shown as the mean of n = 2 technical replicates. **d,** Heat map quantifying *Sen*DRT9 defense activity for the indicated ncRNA mutations, shown as in **c**. **e,** Plaque assay showing loss of *Sen*DRT9 defense activity against T5 phage for substitutions within the putative ncRNA template region. Residues U123–U126 (U_4_) were mutated to each of the indicated nucleotides; EV, empty vector. **f,** Southern blot analysis of total DNA isolated from cells expressing the WT or U_4_>A_4_ MUT ncRNA from **e**, in the absence or presence of T5 phage infection. Duplicate blots were probed with either oligo-dT_40_ (left) or oligo-dA_40_ (right) to detect poly-dA and poly-dT species, respectively. Sizes from a double-stranded DNA (dsDNA) ladder are marked, from the methylene blue-stained membrane after transfer (**Extended Data Fig. 3g**).

To investigate the structural and templating functions of the ncRNA, we scrambled each of the conserved SL regions and assessed the impact of SL perturbation on antiphage defense activity. All SLs were essential for defense against T5 phage except for SL3, whose disruption had a mild effect (**Fig. 2b, Extended Data Fig. 3c**). We then introduced single-nucleotide substitutions within a prominent loop region containing the putative U-rich template. Most mutations in this loop severely impaired defense, suggesting a critical role for this region in ncRNA function (**Fig. 2c**). Strikingly, deletion of a single uridine residue (U124) completely abolished defense, whereas similar perturbations in a secondary U-rich region at the ncRNA’s 3′ end had no effect (**Fig. 2a,d, Extended Data Fig. 3d**). Importantly, RIP-seq experiments on a subset of ncRNA mutants revealed that the observed loss of defense activity was in most cases not attributable to defects in RT-ncRNA binding (**Extended Data Fig. 3e**). Results obtained with the same ncRNA perturbations tested against T2 phage closely mirrored the results with T5 (**Extended Data Fig. 3f**), further supporting the functional importance of ncRNA structural and sequence motifs.

We sought to investigate further the putative templating role of the U-rich loop, and we hypothesized that an uninterrupted stretch of identical nucleotides might be essential for facilitating long homopolymer synthesis. Mutating residues 123–126 (U_4_) to any of the three alternative nucleotides (C_4_, G_4_, or A_4_) abolished phage defense in all cases (**Fig. 2e**), initially suggesting impaired DNA synthesis with these mutants. However, Southern blot analysis of the U_4_>A_4_ mutant revealed a dramatic inversion of the homopolymer synthesis pattern: instead of primarily producing poly-dA with low levels of poly-dT, the A_4_ template primarily generated poly-dT with lower levels of accompanying poly-dA (**Fig. 2f, Extended Data Fig. 3g**). Technical challenges related to the chemical synthesis of guanosine- and cytidine-only oligonucleotides prevented us from testing whether the U_4_>G_4_ and U_4_>C_4_ variants could similarly template complementary homopolymers by Southern blotting.

These results implicate the U-rich loop as an RNA template for complementary DNA (cDNA) synthesis. Like DRT2 systems, which perform rolling circle reverse transcription of a ∼120-nt template to generate concatemeric cDNA products^4,5^, *Sen*DRT9 also spools out concatemeric cDNA products. However, unlike *Kpn*DRT2, *Sen*DRT9 exclusively templates a single nucleotide identity, yielding uninterrupted homopolymers.

### Oligomerization- and protein priming-dependent DNA homopolymer synthesis

We next sought to determine the biochemical and structural basis for RNA-templated homopolymeric DNA synthesis by DRT9. We initially cloned and expressed *Sen*RT as a His_6_-GST fusion protein alongside the ncRNA enriched by RIP-seq (**Extended Data Fig. 4a**). Affinity and size-exclusion chromatography yielded a sample containing the purified RT and an associated RNA species corresponding to the first ∼150 nt of the full-length ncRNA transcript, despite the presence of contaminating proteins (**Extended Data Fig. 4b,c**). Given that His_6_-GST-tagged RT constructs exhibited reduced phage defense activity *in vivo* (**Extended Data Fig. 4d**), we hypothesized that removal of the affinity tag and solubilization domain might be critical for RT activity. Interestingly, when we compared the gel filtration profiles before and after protease cleavage of the N-terminal tag, we observed a marked shift in the retention volume to a monodisperse, higher molecular weight (MW) species, consistent with likely oligomerization (**Extended Data Fig. 4e**). This behavior is reminiscent of oligomerization in AbiK, AbiA, and AbiP2 systems^15,16^, which also exhibit RT self-assembly, though a key distinction is that the *Sen*RT complex includes a ncRNA template.

Incubation of this oligomeric species with [α-^32^P]-dATP, followed by denaturing urea-PAGE analysis, revealed robust and highly processive poly-dA synthesis activity that completely consumed dATP in the reaction, requiring no additional molecular components beyond the RT and ncRNA (**Fig. 3a**). Poly-dA products were 100s to >1000 nt in length and insensitive to the addition of alternative nucleotide substrates (**Fig. 3a, Extended Data Fig. 4f**), corroborating the conclusions from cDIP-seq and Southern blot analyses that DRT9 produces homopolymeric DNA. However, these results also introduced an apparent discrepancy: *in vitro* poly-dA synthesis occurs constitutively, whereas *in vivo* poly-dA synthesis is strictly phage-dependent. This suggested the presence of additional regulatory mechanisms that control DRT9 activation in its native cellular context (see below).

**Figure 3.**
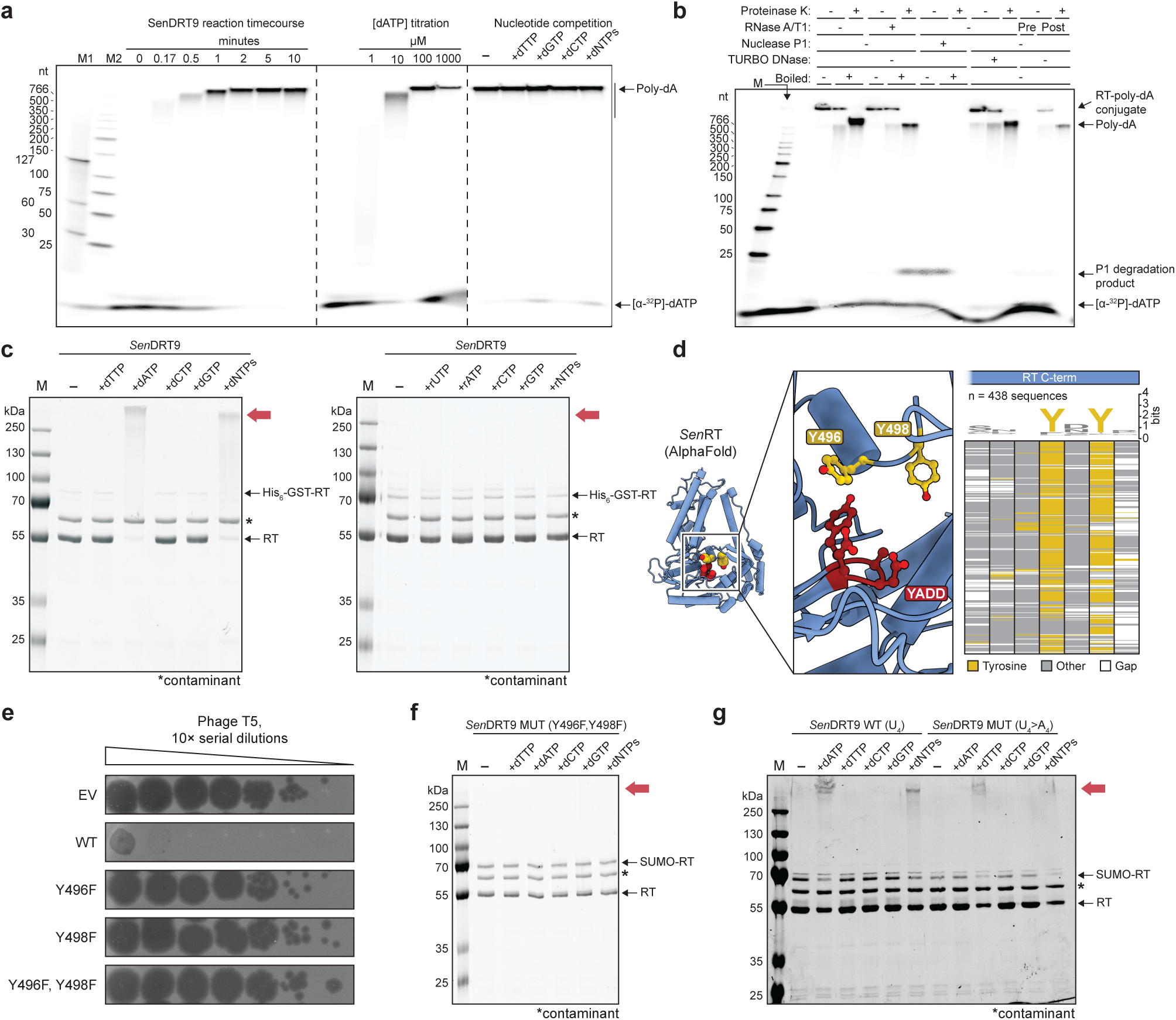
The *Sen*DRT9 RT-ncRNA complex performs protein-primed, RNA-templated DNA homopolymer synthesis. **a,** Denaturing 5% urea-PAGE analysis of DNA polymerization assays that investigated the effect of time (left), dATP concentration (middle), or nucleotide competition (right) on poly-dA synthesis using radiolabeled [α-^32^P]-dATP. All reactions contained 20 nM RT-ncRNA complex and were incubated at 37 °C. *Left*, reactions contained 100 µM dATP; *Middle*, reactions were incubated for 10 min; *Right*, reactions were incubated for 10 min with 100 µM radiolabeled dATP and the indicated, additional unlabeled nucleotides at 100 µM. M1 and M2 denote DNA ladder markers. **b,** Denaturing 5% urea-PAGE analysis of DNA polymerization assays, after reactions were treated with the indicated proteinase or nuclease reagents prior to electrophoretic separation, with or without boiling samples first. All reactions contained 150 nM RT-ncRNA and 100 µM [α-^32^P]-dATP, and were incubated at 37 °C for 10 min prior to proteinase/nuclease addition, with the exception of lane 15, which was pre-treated with RNase A/T1 prior to dATP addition. The prominent mobility shift of the poly-dA product upon proteinase K treatment, irrespective of boiling, suggests the existence of a covalent protein-DNA conjugate. Note that single-strand-specific endonuclease P1 readily degrades poly-dA to small fragments, whereas TURBO DNase, which prefers dsDNA, has little activity on the poly-A product under these conditions. **c,** SDS-PAGE analysis of DNA polymerization assays, in which RT-ncRNA complexes (0.15 µM) were incubated with the indicated dNTPs or rNTPs (0.9 mM) for 60 min at 37 °C, before reactions were quenched and resolved electrophoretically. Red arrows denote smears corresponding to high-molecular weight (MW) protein-DNA conjugates in reactions that contained dATP. Incomplete tag removal during purification accounts for the residual His_6_-GST-RT band; * refers to a purification contaminant; M, protein marker. **d,** AlphaFold 3 structure prediction of a *Sen*RT monomer (left), with magnified view (middle) showing the close proximity of C-terminal residues Y496 and Y498 (yellow) to the YADD active site (red). The multiple sequence alignment (right) highlights the strong conservation of both tyrosine residues. **e,** Plaque assay showing loss of *Sen*DRT9 defense activity against T5 phage for single Y496F and Y498F mutants, as well as a double Y496F,Y498F mutant; EV, empty vector. **f,** SDS-PAGE analysis of DNA polymerization assays as in **c**, but with a Y496F,Y498F RT mutant. **g,** SDS-PAGE analysis of DNA polymerization assays as in **c**, with either WT or a U_4_>A_4_ ncRNA mutant. Mutation of the U_4_ template region to A_4_ completely switches substrate specificity from dATP to dTTP.

Additional observations from analyses of RT reaction products provided insights into the mechanism of poly-dA synthesis. Omitting a proteinase K pre-treatment step noticeably altered the mobility of the poly-dA product, whereas RNase treatment had no effect (**Fig. 3b**). Meanwhile, single-strand-specific nuclease P1 degraded the DNA completely, as expected, whereas TURBO DNase — which preferentially degrades dsDNA — had little effect (**Fig. 3b**). Strikingly, when RT-ncRNA complexes were incubated with various unlabeled dNTP/NTP substrates and then analyzed by SDS-PAGE, we observed a dramatic mobility shift of the RT protein in reactions that contained dATP (**Fig. 3c**). Consistent with these results, gel filtration analysis of the RT-ncRNA complex following dATP incubation revealed complete conversion into a distinct species that exhibited a heightened A_260_/A_280_ ratio, indicating increased nucleic acid content, and ran in the void volume, suggesting a substantial increase in molecular weight (**Extended Data Fig. 5a**). Taken together, these results suggested that poly-dA products were covalently linked to the RT protein itself, potentially via direct priming by a hydroxyl-containing amino acid. Closer inspection of the AlphaFold3-predicted RT monomer structure revealed two highly conserved tyrosine residues within the C-terminal tail, positioned in the immediate vicinity of the active site (**Fig. 3d**). Given the established role of a conserved tyrosine residue in protein-primed DNA synthesis by the AbiK RT^16^, we hypothesized that one or both of these tyrosine residues could similarly serve as primers for reverse transcription by the DRT9 RT.

To test this hypothesis, we individually or simultaneously mutated each conserved tyrosine residue to phenylalanine and assessed the impact on DRT9 activity. Phage defense against T5 was completely abolished when either tyrosine residue was mutated (**Fig. 3e**). In contrast, loss of defense against T2 required substitution of both tyrosines (**Extended Data Fig. 5b**), suggesting that either residue can serve as a potential priming site. RIP-seq experiments in T5-infected cells showed that Tyr>Phe RT mutants exhibited mildly decreased ncRNA binding relative to the WT RT (**Extended Data Fig. 5c**), suggesting that the observed loss of defense may have also resulted in part from weakened RT-ncRNA interactions. Next, we purified the double Tyr>Phe mutant RT-ncRNA complex and found that poly-dA synthesis activity was severely attenuated, as demonstrated by both urea-PAGE analysis of radioactive poly-dA reaction products and SDS-PAGE analysis of RT-DNA conjugates (**Fig. 3f, Extended Data Fig. 5d**). Consistent with the loss of poly-dA synthesis observed *in vitro*, Southern blotting analysis of phage-infected cells expressing Tyr>Phe RT mutants also revealed a strong reduction in poly-dA levels compared to the WT RT (**Extended Data Fig. 5e**). The residual presence of low levels of poly-dA product in both the *in vitro* and *in vivo* experiments with Tyr>Phe RT mutants suggests that additional hydroxyl-containing residues in the RT active site could perhaps be used as alternative DNA synthesis primers, albeit with considerably reduced efficiencies.

Finally, we sought to further assess our model for uridine templating of poly-dA synthesis through biochemical experiments. After purifying RT-ncRNA complexes containing the same U_4_>A_4_ substitution tested in phage defense assays, we performed reverse transcription assays with distinct dNTP substrates. The U_4_>A_4_ mutant generated high-MW protein-DNA conjugates, as assessed by SDS-PAGE, but only in the presence of dTTP (**Fig. 3g**), representing a complete switch in nucleotide substrate specificity. Thus, we conclude that DRT9 systems encode RNA-templated, protein-primed reverse transcriptases that synthesize long, homopolymeric cDNAs covalently linked to specific C-terminal tyrosine residues.

### Cryo-EM structure of the *Sen*DRT9 immune complex

To better understand the molecular and structural basis for this unusual enzymatic behavior, we first determined the structure of the His_6_-GST-tagged RT and ncRNA using cryogenic electron microscopy (cryo-EM) (**Extended Data Fig. 6a–i**). A 3.0 Å-resolution reconstruction enabled unambiguous assignment of 98% of the residues in the RT and 80% of nucleotides in the ncRNA (**Supplementary Table 3**). The RT contains the three characteristic finger, thumb, and palm domains of canonical reverse transcriptases, and an N-terminal extension (residues 1-58) reaches towards the C-terminal thumb domain, turning the conventional right-handed shape of the RT into a triangular architecture (**Fig. 4a, Extended Data Fig. 7a**). The ncRNA forms a complex series of stem loops that interact with every domain of the protein (**Fig. 4b-d**). The 3′ end of the ncRNA curls around the thumb, threading through the center of the triangle and forming extensive, non-base pairing interactions with the 5′ end of the ncRNA (**Fig. 4b**). Notably, the four conserved uridines (123–126) are positioned ∼8 Å from the YADD active site, consistent with a role in templating poly-dA synthesis. However, the C-terminal seven residues, including the two conserved tyrosines (Y496 and Y498), are not ordered.

**Figure 4.**
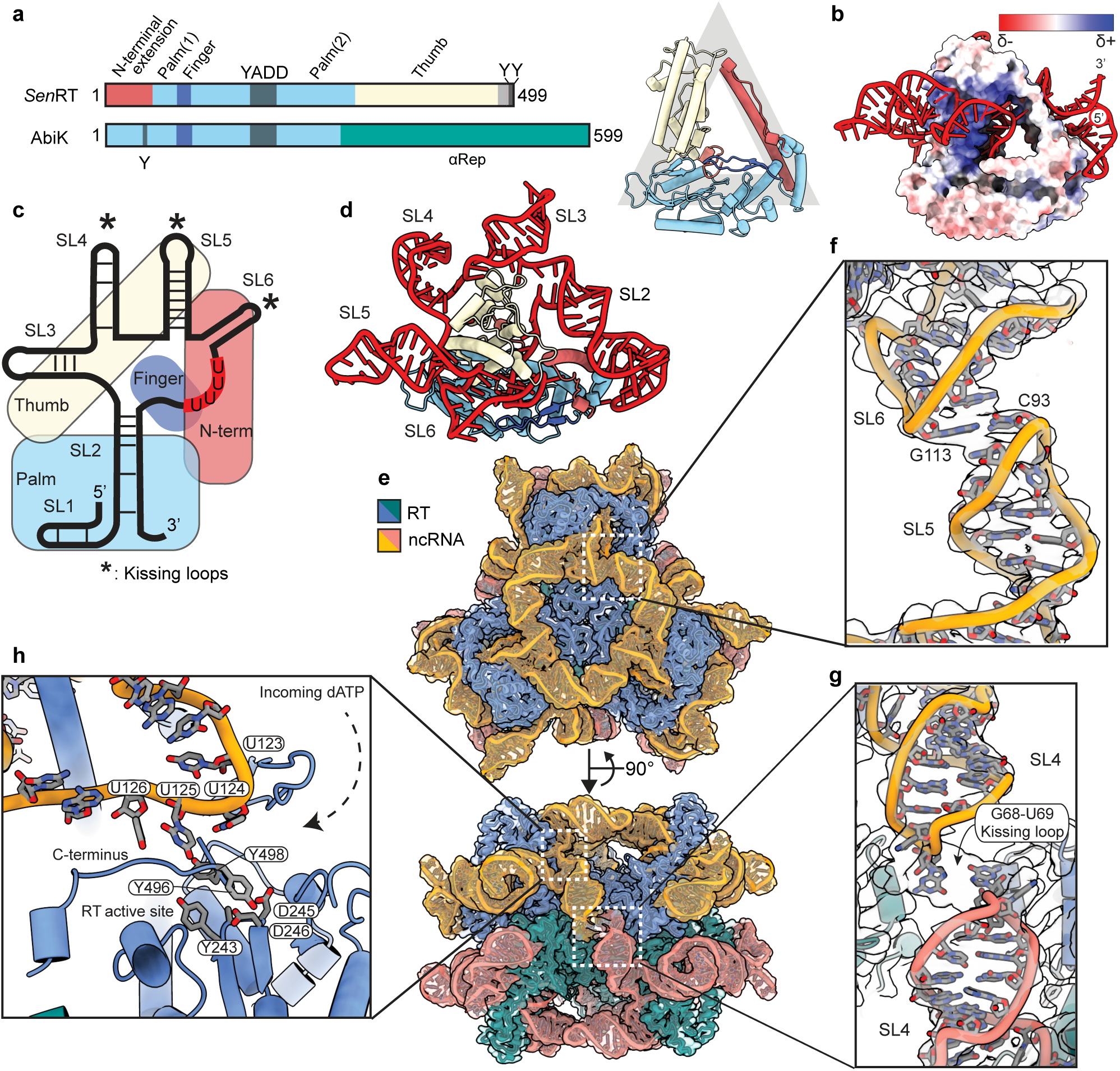
CryoEM structure of the hexameric *Sen*DRT9 RT-ncRNA complex. **a,** Domain architecture (left) of *Sen-* DRT9-encoded RT (PDB: 9NLX, this study) compared to AbiK (PDB: 7R06). The *Sen*RT has an N-terminal extension (red) that forms a triangle-shaped protomer through contacts with the downstream thumb that are bridged by the palm domain (right). **b,** Surface representation of a single RT-ncRNA monomer, colored by relative electrostatic potential, reveals a positively charged tunnel that threads the U_4_ template through the ‘triangle’ of each RT protomer. **c,** Cartoon representation of the ncRNA secondary structure observed in the cryo-EM density, revealing a complex RNA fold composed of several notable features. RNA structures implicated in inter-subunit interactions via kissing loop interactions are highlighted with asterisks. A cartoon representation of the RT domain architecture, colored as in **a**, highlights the path of the RNA across each protein domain. **d,** Structure of a single RT-ncRNA monomer, with RT colored as in **a** and ncRNA stem loops labelled as in **c**, highlighting how each RT protomer is enveloped by its associated ncRNA. **e,** 2.6 Å map and resulting model of the hexameric *Sen*DRT9 RT- ncRNA complex. **f,** Kissing loop interactions between flipped out bases in SL5 (C93) and SL6 (G113) of adjacent ncRNA chains, which form an oligomerization interface above each trimer. **g,** In addition to stabilizing interactions within the trimer, the ncRNA also supports the back-to-back arrangement of trimers via kissing loop interactions between flipped out bases G68 and U69. **h,** A close-up view of the polymerase active site demonstrates how priming tyrosines Y496 and Y498 lie between the YADD(243-246) catalytic motif and templating nucleotides U123-U126, in an orientation poised for homopolymer synthesis.

The RT-ncRNA complex assembles into a C3 trimer stabilized by kissing loop interactions between SL5 (C93) of one molecule and SL6 (G113) of the adjacent subunit. While the structure explains RT-ncRNA interactions and the role of the ncRNA in driving trimer assembly, the affinity-tagged protein is inactive for poly-dA synthesis *in vitro* and non-functional for defense against T5 phage *in vivo* (**Extended Data Fig. 4d,6j**). We therefore hypothesized that removal of the affinity tag would clarify the structural basis for RT-ncRNA function. Following protease cleavage of the affinity tag, we used cryo-EM to determine a 2.6 Å-resolution reconstruction, which revealed a hexameric architecture — a back-to-back dimer of trimers — and ordering of the RT C-terminus (**Fig. 4e, Extended Data Fig. 8, Supplementary Table 3**). The hexameric assembly is reminiscent of AbiK^16^, but the mechanisms of assembly are distinct (**Extended Data Fig. 7b**). Assembly of the AbiK hexamer relies on an alpha-helical repeat domain (i.e., αRep), while assembly of the RT-ncRNA hexamer relies on a combination of RNA-RNA (SL5:SL6 and SL4:SL4), RNA-protein, and protein-protein interactions, resulting in a buried surface area of 3,000 Å^2^ between trimers (**Fig. 4e-g**). While the overall fold of the trimer is nearly identical to equivalent residues in the hexamer, the C-terminal tyrosines (Y496 and Y498) are well ordered in the hexamer and positioned between the active site and the template uridines (123–126), providing further support for the role of the C-terminal tyrosines as primers for DNA synthesis (**Fig. 4h**). Overall, these structural data reveal that the RT-ncRNA complex forms a D3 symmetric hexamer that performs RNA-templated and protein-primed poly-dA synthesis.

### Host and viral factor requirements during DRT9 immunity

To identify viral factors that trigger DRT9 activity, we decided to investigate ‘escaper’ phages harboring mutations that confer resistance to DRT9 immunity. Using a well-established workflow to isolate escaper phages (**Fig. 5a**), we isolated 41 independent T5 phage variants (11 unique genotypes) that evaded *Sen*DRT9, and 1 T5 phage variant and 13 independent Bas37 phage variants (4 unique genotypes) that evaded *Psa*DRT9. Phages that infected DRT9 strains similarly to an empty vector (EV) control were whole-genome sequenced to identify mutations linked to immune evasion (**Extended Data Fig. 9a, Supplementary Table 4**). Sequencing revealed diverse frameshift and missense mutations in *nrdA* (T5, Bas37) and *nrdB* (Bas37), which encode the large and small subunits of viral aerobic ribonucleoside-disphosphate reductase (RNR) complexes that catalyze ribonucleotide to deoxyribonucleotide conversion. This immediately implicated nucleotide availability as a potential vulnerability of DRT9 immunity and/ or phage infection, particularly given the rapid consumption of dATP by RT-ncRNA complexes *in vitro*. Additionally, previous studies have suggested that T5 replication is particularly sensitive to dTTP levels in infected cells^18^, suggesting that DRT9 might restrict T5 genome replication by depleting nucleotide levels. To test this hypothesis, we measured nucleotide levels in uninfected and T5 phage-infected cells and found that in infected cells, *Sen*DRT9 expression diminished adenosine nucleotide levels compared to EV controls (**Extended Data Fig. 9b**). However, supplementing the growth medium with exogenous nucleosides had no discernible effect on DRT9 immunity or phage replication (**Extended Data Fig. 9c**), suggesting a more complex regulatory mechanism.

**Figure 5.**
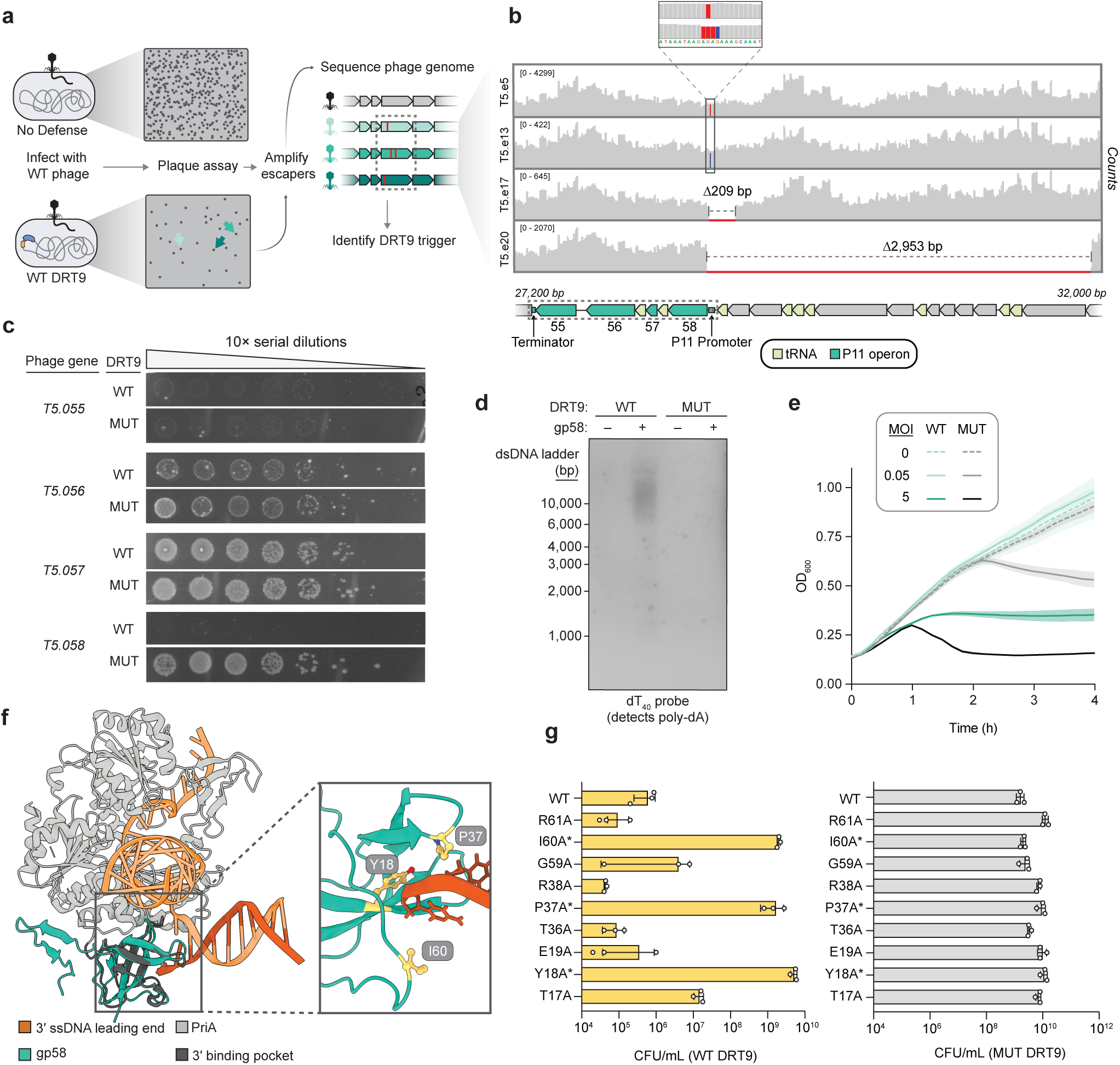
Identification of a viral trigger that activates *Sen*DRT9-mediated abortive infection. **a,** Schematic of workflow to isolate phage variants that escape detection and/or elimination by a *Sen*DRT9 immune response. Individual plaques escaping defense are isolated and sequenced to identify mutations affecting candidate trigger genes. **b,** Representative coverage tracks from whole-genome sequencing of T5 ‘escaper’ phage variants that bypass *Sen*DRT9 immunity, shown above the corresponding annotated genomic locus. All mutation classes shown (point mutations and larger deletions) perturb the putative P11 promoter driving expression of four protein-coding genes. **c,** Colony formation assays to determine cell viability upon co-expression of WT or RT-inactive (MUT) *Sen*DRT9 with candidate phage genes regulated by the P11 promoter, identified in **b**. The gene product of *T5.058* (gp58) prevents cell growth exclusively in WT *Sen*DRT9 cells, whereas gp55 causes generic toxicity regardless of *Sen*DRT9 activity; the remaining candidates exhibit no phenotype. **d,** Southern blot analysis of total DNA isolated from cells expressing WT or MUT *Sen*DRT9, with or without gp58 induction. DNA was probed with oligo-dT_40_ to detect poly-dA species. Sizes from a double-stranded DNA (dsDNA) ladder are marked, from the methylene blue-stained membrane after transfer (**Extended Data Fig. 9d**). **e,** Growth curves of cells expressing WT or MUT *Sen*DRT9, in the presence or absence of T5 phage at the indicated multiplicity of infection (MOI). Shaded regions indicate the SD across n = 3 independent biological replicates. **f,** Predicted AlphaFold 3 structure of T5 gp58 (teal) superimposed onto the structure of the *E. coli* replication restart helicase PriA (grey) bound to a replication fork DNA substrate (orange; PDB ID: 8FAK). The magnified inset highlights gp58 residues that overlap PriA residues implicated in binding to DNA 3′ ends^22^. RMSD = 0.95 Å over 47 Cα atoms between gp58 and PriA. **g,** Bar graphs quantifying cell viability in colony forming units (CFU) upon co-expression of the indicated gp58 alanine substitution variants with *Sen*DRT9 WT (left) or MUT (right). Asterisks (*) indicate amino acid residues predicted to interact with the DNA substrate 3′ end, as highlighted in **f**. Data are shown as mean ± s.d. for n = 3 technical replicates.

A larger set of T5 escaper mutations were clustered within a well-defined genomic region, consisting primarily of deletions spanning 200–15,000 bp (**Supplementary Table 4**). This tRNA-rich region, known as del-1, is a hotspot for large genomic deletions due to repeat-associated recombination^19,20^. However, the discovery of additional escaper genotypes containing only point mutations within an intergenic region redirected our attention to the P11 promoter, which drives expression of a six-gene operon that includes two tRNAs (**Fig. 5b**). We hypothesized that one or more genes within this operon might activate the DRT9 immune response, and cloned expression plasmids for each of the four protein-coding genes to test the effects of DRT9 and phage gene co-expression. Strikingly, a single gene, *T5.058*, prevented cell growth when co-expressed with WT *Sen*DRT9 but not the RT-inactive mutant, suggesting its potential role in activating poly-dA synthesis (**Fig. 5c**). This was supported by Southern blotting analysis, which showed accumulation of poly-dA products in cells co-expressing *T5.058* with WT *Sen*DRT9 (**Fig. 5d, Extended Data Fig. 9d**), albeit at lower levels than those induced by phage infection. Consistent with the growth defect observed upon co-expression of *Sen*DRT9 and *T5.058*, phage infection assays in liquid culture revealed that cells expressing *Sen*DRT9 exhibited a growth arrest phenotype upon immune system activation by a high multiplicity of T5 infection (**Fig. 5e**). These results indicate that the overall outcome of phage-triggered DRT9 activation is growth arrest and/or cell death, and strongly implicate gp58, the *T5.058* gene product, as a necessary and sufficient T5 phage factor for triggering DRT9-mediated abortive infection.

We predicted the structure of gp58, an uncharacterized protein, and identified a high degree of structural similarity to the 3′ DNA-binding domain (3′BD) of PriA (**Fig. 5f**), a primosomal protein involved in DNA replication restart at stalled or damaged replication forks^21^. Co-immunoprecipitation experiments with FLAG-RT and V5-gp58 revealed a direct interaction in cells expressing the WT *Sen*DRT9 system, but not the RT-inactive mutant (**Extended Data Fig. 9e**), implicating gp58 in RT- and/or poly-dA-dependent binding. To further probe its role, we used the gp58 structure prediction, alongside previous studies of DNA binding by PriA^22^, to design gp58 variants predicted to disrupt DNA interactions. Mutating gp58 in this manner restored normal growth in cells co-expressing gp58 and *Sen*DRT9, indicating failed triggering of abortive infection (**Fig. 5g**). Thus, we conclude that gp58 likely recognizes the DNA homopolymer synthesized by *Sen*DRT9, potentially through direct binding to its 3′ end.

Our finding that DRT9 activity is triggered by a phage protein predicted to bind DNA 3′ ends dovetailed with a separate insight that arose from literature review. A comprehensive study of *E. coli* gene essentiality determined that *sbcB*, which encodes the 3′-to-5′ exodeoxyribonuclease I (ExoI), is an essential gene in strains expressing UG17 (an RT system related to DRT9), suggesting that ExoI mitigates the otherwise toxic effects of UG17^23^. Strikingly, we were unable to obtain transformants of a *ΔsbcB* strain with WT *Sen*DRT9, whereas the RT-inactive mutant was well-tolerated (**Fig. 6a**), revealing a similar conditional essentiality relationship. This discovery suggested a model in which ExoI constitutively degrades DRT9-synthesized poly-dA products in uninfected cells, but is out-competed for cDNA 3′ end recognition by gp58 during T5 phage infection, leading to poly-dA accumulation and abortive infection. Importantly, this mechanism would resolve the apparent discrepancy between our *in vivo* observation that poly-dA synthesis occurs only after phage infection, and our *in vitro* observation that poly-dA synthesis activity occurs constitutively, with no molecular requirements other than the RT and ncRNA. In excellent agreement with this model, recombinant ExoI potently degraded DRT9-derived poly-dA products, but this activity was directly inhibited by gp58 in a concentration-dependent manner, whereas the same was not true for nuclease P1 (**Fig. 6b, Extended Data Fig. 9f**).

**Figure 6.**
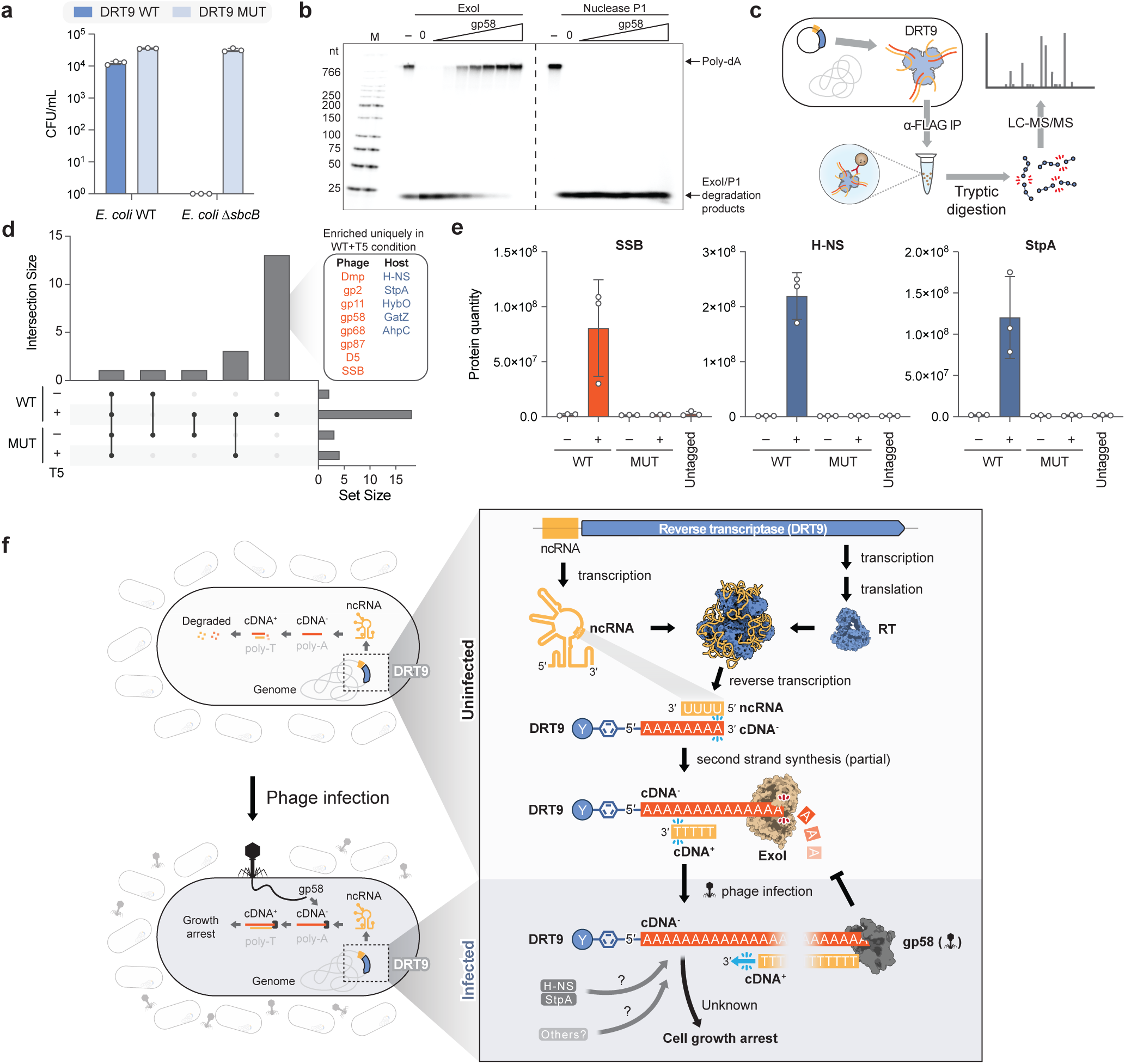
Host regulation of and response to poly-dA synthesis by DRT9 immune systems. **a,** Bar graph quantifying cell viability in colony forming units (CFU) upon transformation of WT or *ΔsbcB E. coli* with either WT or RT-inactive (MUT) *Sen*DRT9. The inability to propagate WT *Sen*DRT9 in a Δ*sbcB* background suggests that the *sbcB* gene product, Exodeoxyribonuclease I (ExoI), neutralizes the otherwise toxic properties of DRT9 synthesis products. Data are shown as mean ± s.d. for n = 3 technical replicates. **b,** Denaturing 5% urea-PAGE analysis of DNA polymerization assays, after reactions were treated with ExoI (left) or DNase P1 (right) in the presence of increasing concentrations of T5 gp58. All reactions contained 20 nM RT-ncRNA and 100 µM [α-32P]-dATP, and were incubated at 37 °C for 10 min prior to incubation with 0.015 Units/µL ExoI or 2 Units/µL nuclease P1 alongside gp58 ranging from 0.1–10 µM. Nuclease P1, which is an endonuclease, is not inhibited by gp58. **c,** Schematic of co-immunoprecipitation followed by mass spectrometry (co-IP MS) to identify protein interactors of *Sen*DRT9; proteins that interact with RT-mediated DNA synthesis products will be enriched alongside direct interactors with the RT-ncRNA complex. **d,** UpSet plot showing the number of overlapping and unique protein interactors identified from co-IP MS experiments with either WT or MUT *Sen*DRT9 in the absence or presence of T5 phage infection. The size of each significantly enriched protein set is displayed on the bar graph to the right, while the size of set overlaps is displayed on the bar graph above. Significantly enriched proteins were identified as those exhibiting >20-fold enrichment relative to the control IP, with a false discovery rate (FDR) < 0.05. Phage and host proteins enriched uniquely in the WT + T5 condition are shown at right, which notably includes the T5 gp58 trigger protein and host nucleoid-associated proteins (NAPs) such as H-NS and StpA, known to preferentially bind AT-rich DNA. **e,** Bar graph showing normalized protein intensity values for T5 SSB and *E. coli* H-NS and StpA across the indicated co-IP MS experiments from **d**, including an untagged (no FLAG) control. Data are shown as mean ± s.d. for n = 3 independent biological replicates. **f,** Model for the antiphage defense mechanism of DRT9 systems. The RT and ncRNA form a hexamer, leading to constitutive, protein-primed synthesis of the poly-dA cDNA by the U-rich template region of the ncRNA, and partial second-strand synthesis of poly-dT. Host ExoI normally degrades poly-dA products in the cell, but phage infection leads to trigger expression (gp58 in T5) that competes for 3′ end binding, thereby stabilizing poly-dA products and causing their accumulation. *Sen*DRT9 poly-dA products are bound by additional viral and host factors in the cell (H-NS and StpA), leading to cell growth arrest through an unknown mechanism.

Finally, we repeated immunoprecipitation experiments using the FLAG-tagged RT in T5 phage-infected cells, followed by unbiased proteomics analysis, to identify additional viral or host factors involved in DRT9 immunity (**Fig. 6c**). As expected, gp58 was significantly enriched in cells expressing WT *Sen-* DRT9 but not the RT-inactive mutant, corroborating our earlier findings by Western blotting. Additional T5 proteins implicated in DNA binding and genome replication were also uniquely enriched by the WT RT, including T5 single-stranded DNA binding protein (SSB) (**Fig. 6d,e)**. Intriguingly, immunoprecipitation of the WT RT also strongly enriched two host-encoded nucleoid-associated proteins, H-NS and StpA (**Fig. 6d,e**). Both proteins preferentially bind AT-rich DNA regions and are key regulators of transcription and nucleoid architecture^24^, providing an elegant link between DRT9-catalyzed cDNA synthesis and associated host factors.

## DISCUSSION

Our work uncovers an unprecedented mechanism of antiviral immunity mediated by DRT9 defense systems, in which a noncoding RNA (ncRNA) templates the synthesis of long, kilobase-length homopolymers of deoxyadenylate (poly-dA) to drive abortive infection (**Fig. 6f**). Biochemical assays demonstrate that poly-dA synthesis is dependent on only the reverse transcriptase (RT) and ncRNA, while a conditional essentiality relationship between DRT9 and *sbcB* revealed that the *sbcB* gene product ExoI constitutively degrades poly-dA products in uninfected cells, suppressing its toxic effects. During infection by T5 phage, expression of phage gp58 — a predicted DNA binding protein that blocks ExoI access to the poly-dA product — leads to poly-dA accumulation in cells and growth arrest. Importantly, our finding that gp58 alone is sufficient to induce toxicity in a DRT9-expressing strain argues against poly-dA synthesis acting to specifically antagonize phage infection. Instead, we propose that poly-dA accumulation functions as the effector arm of DRT9 immunity by driving a phage-extrinsic abortive infection mechanism.

The mechanism by which poly-dA accumulation leads to abortive infection remains elusive. *In vitro* reverse transcription reactions with *Sen*DRT9 rapidly and quantitatively consumed dATP, leading us to initially suspect that nucleotide depletion might inhibit both host and phage genome replication by starving the cell of adenosine, thereby explaining both DRT9 immunity and abortive infection. If true, this mechanism would provide an elegant counterpart to other phage defense systems that similarly deplete deoxyribonucleotides^8,9^. However, while nucleotide metabolomics measurements revealed a clear DRT9-dependent decrease in cellular dATP levels in infected cells, supplementation of deoxyadenosine nucleosides failed to suppress the immune response (**Extended Data Fig. 9b,c**). An alternative hypothesis arises when considering the observation that DRT9 systems sometimes associate with SLATT domains (**Extended Data Fig. 1a**), which contain transmembrane (TM) helices predicted to form membrane-embedded pores that function as toxin effectors^25^. The recurring presence of SLATT and TM domains in other families of Unknown Group reverse transcriptase systems^3^, alongside recent reports implicating oligonucleotide-activated pore opening in abortive infection responses of other phage defense systems^26,27^, suggests that nucleic acid-mediated regulation of membrane proteins may be a broader theme in antiviral immunity. However, whether the poly-dA products of *Sen*DRT9 immunity similarly engage membrane proteins in *E. coli* requires future investigation.

The unusual mechanism of RNA-templated homopolymer synthesis reported here sets DRT9 reverse transcriptase enzymes apart from all other known polymerases. While nucleotidyltransferase (NTase) enzymes that polymerize RNA substrates — such as poly(A) and poly(U) polymerase — also produce nucleic acid homopolymers, they do so without a template, relying instead on protein-mediated nucleotide selection within the active site^28^. Similarly, terminal deoxynucleotidyl transferase (TdT) can add homopolymeric stretches of DNA when a given nucleotide is in excess, yet lacks the specificity conferred by a guiding nucleic acid sequence^29^. DRT9 enzymes are, to our knowledge, the only polymerases that harness an RNA template and reverse transcriptase domain to direct the synthesis of DNA homopolymers, yielding kilobase-length products in a highly processive manner. Even more strikingly, these homopolymers remain tethered to the enzyme itself as covalent conjugates to the C-terminal tail, via a protein priming mechanism dependent on conserved tyrosine residues.

The structural architecture of DRT9 RT-ncRNA complexes is equally distinctive, especially in comparison to DRT2-family enzymes, whose members occupy neighboring positions on the phylogenetic tree of UG RTs (**Fig. 1a**). While both RTs share a similar domain architecture and engage ncRNA substrates with comparable secondary structures, their assembly modes are strikingly different. The DRT2 ribonucleoprotein (RNP) complex behaves as a functional monomer^5^, whereas the DRT9 RNP adopts a hexameric architecture comprising a dimer of trimers. This arrangement echoes the hexameric and trimeric assemblies of two other UG-family reverse transcriptases, AbiK and Abi-P2^16^, but is uniquely stabilized by an intricate network of RNA-dependent interactions that form a tightly interlinked cage to encapsulate the enzyme subunits. The proximity of the U-rich template region and C-terminal tail to the catalytic pocket provides a structural framework for understanding the mechanism of poly-dA synthesis. Indeed, the proposed mechanism by which DRT9 RTs iteratively engage the same template region for repetitive addition of identical nucleotides is highly reminiscent of repeat addition processivity (RAP), a hallmark feature of telomerase reverse transcription activity during telomeric DNA elongation^30^. Future studies will be needed to determine the precise number of uridines required for templated deoxyadenosine incorporation, how the nascent cDNA is realigned and reset within the active site, and intriguingly, how the kilobase-sized homopolymers extrude from the body of the hexameric RNP while remaining covalently tethered at the 5′ end and engaged within the active site at the 3′ end.

As with DRT2, discoveries on DRT9 emerged from challenging conventional expectations about RT enzymes and their cDNA synthesis products. In both cases, a departure from standard approaches in high-throughput sequencing data analysis allowed us to identify unexpected cDNA products. Notably, our initial interest in DRT9 stemmed from the hypothesis that its properties would closely resemble DRT2, given their close phylogenetic relationship, but our findings here illuminate a dramatically different mechanism of antiviral immunity. As such, we anticipate that further exploration of reverse transcriptase diversity will continue to uncover unexpected DNA synthesis mechanisms, and that the full scope of nucleic acid functions in antiviral defense is only beginning to come into view.

## METHODS

### Plasmid and *E. coli* strain construction

All strains and plasmids used in this study are described in **Supplementary Tables 5–6**, respectively. DRT9 operons, including their native upstream and down-stream flanking sequences, were chemically synthesized (Twist Bioscience or GenScript) and cloned into a pACYC184 plasmid backbone by Gibson Assembly. Derivative plasmids were cloned using a variety of methods, including around-the-horn PCR, restriction digestion-ligation, and Golden Gate Assembly. All plasmids were cloned and propagated in *E. coli* strain NEB Turbo (sSL0410). Clones were verified by Sanger sequencing or whole plasmid sequencing. Substitution and insertion mutations to the *Sen*DRT9 ncRNA are numbered relative to the first nucleotide of the ncRNA, with insertion numbering referencing the nucleotide immediately upstream of the inserted bases (for instance, A109U indicates mutation of A at position 109 to U, and 124insU indicates insertion of U downstream of position 124). Experiments were performed in *E. coli* K-12 strain MG1655 unless otherwise indicated. The Δ*sbcB* strain from the Keio collection^31^, which harbors a single-gene deletion in *E. coli* K-12 strain BW25113, was a gift from S. Tavazoie.

### Phylogenetic analyses

A BLASTp search of a local copy of the NCBI NR database (downloaded April 4, 2023) was queried with 124 DRT9 (i.e., UG28) sequences identified by Mestre et al.^3^ (-evalue 0.01 -max_target_seqs 1,000,000), resulting in the identification of 438 unique DRT9 homologs. These proteins were aligned with MAFFT^32^ (LINSI option) and a phylogenetic tree was constructed from the resulting alignment with FastTree [-wag -gamma options]^33^. Genomes encoding these DRT9 homologs were then downloaded using NCBI’s Batch-Entrez function, and loci comprising *drt9* plus 10 kbp of upstream and downstream nucleotide sequence were extracted. Next, to identify DRT9 systems that encode a SLATT domain-containing gene, we predicted ORFs in each DRT9 locus using Eggnogg Mapper^34^ and used HMMER^35^ to search the resulting ORFs with a previously built hidden Markov model (PFAM: PF18160). Finally, ncRNA associations were annotated using the CMsearch function of Infernal^36^ to search DRT9 loci with a CM built from DRT9-associated ncRNAs described below.

### Sequence identity matrices

Sequence identity between selected DRT9 homologs was analyzed by performing MAFFT alignment of RT amino acid sequences using default settings. Accession numbers for RT proteins are listed in **Supplementary Table 2**.

### Phage propagation and plaque assays

Phages T2, T5, and λ-vir were gifts from Michael Laub. BASEL collection phages were gifts from Alexander Harms. All phages were propagated in liquid culture by picking single plaques into LB media, supplemented with 5 mM CaCl_2_ and 5 mM MgSO_4_, containing *E. coli* MG1655 cells diluted 1:100 from overnight cultures. After 4–5 hours of incubation with shaking at 37 °C, chloroform was added to a final concentration of 5% to lyse residual bacteria, and lysates were centrifuged at 4,000 x *g* for 10 min to pellet cell debris. Supernatants were passed through a sterile 0.22 µm filter and stored at 4 °C.

Small-drop plaque assays were performed as previously described^4^. Briefly, *E. coli* K-12 strain MG1655 (sSL0810) (see **Supplementary Table 5** for strain descriptions and genotypes) was transformed with the indicated plasmids (see **Supplementary Table 6** for plasmid descriptions and sequences). Transformants were inoculated in liquid LB media containing the appropriate antibiotic and grown overnight at 37 °C with shaking. The next day, overnight cultures were mixed with molten soft agar (0.5% agar in LB media supplemented with 5 mM CaCl_2_ and 5 mM MgSO_4_) at 45 °C and poured over solid bottom agar (1.5% agar in LB media containing the appropriate antibiotic) in a Petri dish. 10× serial dilutions of phage in LB media were spotted onto the surface of the soft agar lawn. Plates were incubated at 37 °C for 8–16 hours to allow plaque formation. Plaque forming units (PFU) mL^−1^ were calculated using the following formula: . Phage defense activity was assessed by calculating the fold reduction in efficiency of plating (EOP), which was determined by dividing the PFU mL^−1^ obtained on a lawn of empty vector (EV)-transformed control cells by the PFU mL^−1^ obtained on a lawn of defense system-expressing cells.

### AlphaFold structural modeling

Protein sequences were submitted to the AlphaFold 3 webserver^37^ and the resulting structural predictions were rendered with ChimeraX^38^.

### RNA and cDNA immunoprecipitation and sequencing (RIP-seq and cDIP-seq)

RIP-seq and cDIP-seq were performed as previously described^4^. *E. coli* K-12 strain MG1655 (sSL0810) cells were transformed with plasmids encoding N-terminally 3xFLAG-tagged DRT9 systems with a WT or mutant (YAAA) RT active site (see **Supplementary Table 6** for plasmid sequences). Individual colonies for each replicate experiment were inoculated in 40 mL liquid LB and grown at 37 °C to OD_600_ of 0.3-0.4. For experiments performed in the absence or presence of phage infection, cultures were split in half and phage T5 was added to one half at a multiplicity of infection (MOI) of 5. Cultures were further incubated at 37 °C with shaking for 45 min before harvesting. For experiments performed only in uninfected cells, 20 mL cultures were grown to OD_600_ of 0.5 and harvested. Cells were collected by centrifugation at 4,000 x *g* for 10 min at 4 °C and then washed with 5 mL of cold TBS (20 mM Tris-HCl pH 7.5 at 25 °C, 150 mM NaCl). Cells were centrifuged again at 4,000 x *g* for 5 min at 4 °C. Pellets were washed with 1 mL of cold TBS and centrifuged at 10,000 x *g* for 5 min at 4 °C. After supernatants were removed, pellets were flash-frozen in liquid nitrogen and stored at –80 °C.

To prepare antibody–bead complexes for immunoprecipitation, Dynabeads Protein G (Thermo Fisher Scientific) were washed 3× in 1 mL IP Lysis Buffer (20 mM Tris-HCl pH 7.5 at 25 °C, 150 mM KCl, 1 mM MgCl_2_, 0.2% Triton X-100), resuspended in 1 mL IP Lysis Buffer, combined with anti-FLAG antibody (Sigma-Aldrich, F3165), and rotated for > 3 hours at 4 °C. 60 µL of beads and 20 µL of antibody mix were prepared per sample. Antibody-bead complexes were washed 2× to remove unbound antibodies and resuspended in IP Lysis Buffer to a final volume of 60 µL per sample.

Flash-frozen pellets were thawed on ice and resuspended in 1.2 mL IP Lysis Buffer supplemented with 1× Complete Protease Inhibitor Cocktail (Roche) and 0.1 U µL−1 SUPERase•In RNase Inhibitor (Thermo Fisher Scientific). Cells were lysed by sonication and lysates were cleared by centrifugation at 21,000 x *g* for 15 min at 4 °C. Supernatants were transferred to new tubes and two 10 µL aliquots of each sample (one for RIP-seq and one for cDIP-seq) were set aside and stored at –80 °C as “input” controls. The remainder of each sample was combined with 60 μL of antibody-bead complex and rotated overnight at 4 °C. The next day, each sample was washed 3× with 1 mL ice-cold IP Wash Buffer (20 mM Tris-HCl pH 7.5 at 25 °C, 150 mM KCl, 1 mM MgCl_2_). During the final wash, each sample was split into two 500 µL volumes for downstream RIP or cDIP processing.

RIP elution was performed by removing supernatants and resuspending beads in 750 µL TRIzol (Thermo Fisher Scientific). After 5 min incubation at RT, supernatants containing eluted RNA were transferred to new tubes. Samples were then mixed with 150 μL chloroform, incubated at RT for 2 min, and centrifuged at 12,000 x *g* for 15 min at 4 °C. RNA was isolated from the upper aqueous phase using the RNA Clean & Concentrator-5 kit (Zymo Research). RNA from input samples was isolated in the same manner using TRIzol and column purification. cDIP elution was performed by removing supernatants, resuspending beads in 90 µL IP Wash Buffer, and treating samples with 5 µg RNase A (Thermo Fisher Scientific) and 25 µg Proteinase K (Thermo Fisher Scientific). Supernatants containing eluted DNA were transferred to new tubes. DNA was isolated using the Monarch Spin PCR and DNA Cleanup kit (NEB), following the Oligonucleotide Cleanup protocol. Input samples were also treated with RNase A and Proteinase K prior to DNA isolation by column purification.

For RIP-seq library preparation, RNA was diluted in NEBuffer 2 and heated at 92 °C for 2 min for fragmentation by alkaline hydrolysis. Samples were then treated with TURBO DNase (Thermo Fisher Scientific), RppH (NEB), and T4 PNK (NEB) to remove DNA and prepare RNA ends for adapter ligation. RNA was purified using the Zymo RNA Clean & Concentrator-5 kit. Adapter ligation, reverse transcription, and indexing PCR were performed using the NEBNext Small RNA Library Prep kit. Libraries were sequenced on an Element AVITI in paired-end mode with 75 or 150 cycles per end.

For cDIP-seq library preparation, samples were heated at 95 °C for 2 min and then immediately placed on ice to denature DNA. Adapter ligation and conversion of ssDNA to dsDNA were performed using the xGen ssDNA & Low-Input DNA Library Prep Kit (IDT). Indexing PCR was performed using NEBNext Ultra II Q5 Master Mix. Libraries were sequenced on an Element AVITI in paired-end mode with 75 or 150 cycles per end.

### RIP-seq analysis

RIP-seq datasets were processed as previously described^4^. Cutadapt^39^ (v4.2) was used to remove adapter sequences, trim low-quality ends from reads, and filter out reads shorter than 15 bp. Reads were mapped to reference files containing the MG1655 genome (NC_000913.3), relevant plasmid sequence, and T5 (NC_005859.1) genome, using bwa-mem2^40^ (v2.2.1) with default parameters. SAMtools^41^ (v1.17) was used to sort and index alignments. Coverage tracks for top and bottom strand alignments were generated using bamCoverage^42^ (v3.5.1) with normalization for sequencing depth (based on the total number of reads passing initial trimming and length filtering). Coverage tracks were visualized in IGV^43^.

For transcriptome-wide analysis of RNAs enriched by RIP-seq, aligned reads were assigned to annotated transcriptome features using featureCounts^44^ (v2.0.2) with -s 1 for strandedness. Counts matrices were analyzed using DESeq2^45^ to calculate the enrichment (fold-change) and false discovery rate for each transcript compared between input and IP samples. Transcriptome-wide comparisons were visualized as MA plots, using ggplot2 to plot “baseMean” (mean normalized counts across all conditions) against log_2_(enrichment). All comparisons included three independent biological replicates.

### cDIP-seq analysis

Adapter trimming, quality trimming, and length filtering of cDIP-seq and corresponding input (total DNA) datasets were performed as described above for RIP-seq experiments. Trimmed and filtered reads were mapped to combined reference files, sorted, indexed, and plotted onto coverage tracks as described above. For transcriptome-wide analysis of cDIP-seq data, alignments over annotated transcriptome features were counted using featureCounts with -s 2 for strandedness. Counts matrices were processed by DESeq2 and plotted as described above. The *lacZ* and *racR* loci were masked from this analysis, as they were previously found to show artifactual enrichment by cDIP-seq^4^. All transcriptome-wide comparisons were performed using three independent biological replicates.

### Unmapped read analysis

For analysis of unmapped reads in cDIP-seq and total RNA-seq datasets (using the input controls from RIP-seq experiments), reads were first stringently trimmed and filtered to remove all residual adapter sequences. Cutadapt was used to process reads, adapting a previously described method for tiling across the 3′ adapter^46^, with error, overlap, quality, and length thresholds set at -e 0.2 -O 10 -q 30 -m 45. The fraction of unmapped reads for each sample was calculated using SAMtools flagstat. Unmapped reads were then extracted for downstream analysis using SAMtools fasta. Motif analysis was performed using MEME^47^ (v5.5.7) in differential enrichment mode with minimum width 45 (-minw 45) and allowing any number of motif repetitions (-mod anr), using unmapped reads from cDIP-seq of the catalytically inactive RT (YAAA) as the control sequence set. Quantification of homopolymer-containing reads was performed using countPattern from the Biostrings package (v2.70.3) in R. Reads were analyzed for the presence of at least 25 consecutive A, T, G, or C bases, or for the presence of A_10_T_10_ or T_10_A_10_ sequences, allowing 1 mismatch. Reads containing homopolymers were counted and normalized according to the total number of filtered reads for each sample.

### Oligonucleotide spike-in sequencing

To assess potential biases in library preparation and sequencing of poly-dA and poly-dT, 100-nt single-stranded DNA homopolymers (oligo-dA and oligo-dT) were chemically synthesized (IDT) and analyzed by next-generation sequencing. 250 pmol oligo-dA and oligo-dT were each 5′-phosphorylated by treatment with T4 PNK (NEB) for 30 min at 37 °C. Reactions were cleaned up using the Monarch Spin PCR and DNA Cleanup Kit (NEB) and eluted in 20 µL water. Phosphorylated oligos were added to ∼5 pmol total DNA — extracted from *E. coli* K-12 strain MG1655 cells expressing catalytically inactive *Sen*DRT9 — in various amounts (50 fmol oligo-dA only, 50 fmol oligo-dT only, 50 fmol each of oligo-dA and oligo-dT, or 500 fmol of oligo-dA and 50 fmol of oligo-dT). Spiked-in DNA samples were subjected to the same library preparation and sequencing workflow as described above for cDIP-seq.

### Southern blotting

DNA samples were collected from either uninfected or phage T5 infected cells. Cultures from a single colony were grown in 20 mL liquid culture until OD_600_ reached 0.3. For experiments performed in presence of phage infection, phage T5 was added at a multiplicity of infection (MOI) of 5. For uninfected cells, an equal volume of LB was added. Cultures were further incubated at 37 °C with shaking for 60 min before harvesting. For experiments containing the gp58 trigger, cells were grown in 20 ml LB supplemented with chloramphenicol (25 µg/mL), spectinomycin (100 µg/mL) and 2% glucose until they reached OD_600_ of 0.3. Cells were pelleted by centrifugation at 4,000 x *g* for 5 min, washed with 10 mL LB and resuspended in 20 mL LB with chloramphenicol (25 µg/mL), spectinomycin (100 µg/mL) and 0.2% L-arabinose to induce gp58 expression. Cultures were incubated at 37 °C with shaking for 45 min before harvesting. Cells were collected by centrifugation at 4,000 x *g* for 10 min at 4 °C and then washed with 5 mL of cold TBS. Cells were centrifuged again at 4,000 x *g* for 5 min at 4 °C. Pellets were washed with 1 mL of cold TBS and centrifuged at 10,000 x *g* for 5 min at 4 °C. Afterwards supernatants were removed and pellets were stored at –20 °C.

Genomic DNA samples were extracted using the Wizard Genomic DNA Purification Kit (Promega). 3 µg of gDNA were loaded on an agarose gel (0.8% agarose, 1× TAE Buffer) and run for 2 hours at 90 V. The gel was soaked for 10 min in Denaturation Buffer (0.5 M NaOh, 1.5 M NaCl), and DNA was transferred to a Hybond-N+ membrane (GE Healthcare) by upward capillary transfer in Alkaline Transfer Buffer (0.25 M NaOH, 1.5 M NaCl). The next day, DNA was denatured for probe hybridization by incubating the membrane in 0.4 M NaOH for 10 min, then rinsed in Na_2_HPO_4_ (0.5 M, pH 7.3) and DI water. DNA transfer was confirmed by staining the membrane with methylene blue and imaging with an Amersham Imager 600 (GE Healthcare). The membrane was pre-hybridized in ULTRAhyb-Oligo Buffer (Thermo Fisher Scientific) for 1 hour at 42 °C. A biotinylated oligonucleotide probe of poly-dT_40_ or poly-dA_40_ was added to the Hybridization Buffer at a final concentration of 5 nM and hybridization was performed overnight at 42 °C. The next day, the membrane was washed twice with Southern Blot Wash Buffer I (2× SSC with 0.1% SDS) and twice with Southern Blot Wash Buffer II (0.1× SSC with 0.1% SDS). The membrane was developed using the Chemiluminescent Nucleic Acid Detection Module Kit (Thermo Fisher Scientific) and imaged with an Amersham Imager 600 (GE Healthcare). Probe sequences are provided in **Supplementary Table 7**.

### ncRNA covariance modeling

Related homologs of DRT9 were identified using the amino acid sequence of a *Salmonella enterica* homolog (*Sen*DRT9, EAO1508900.1) as the seed query in a BLASTp search on the NR protein database (max target sequences = 100). Nucleotide sequences 1 kb upstream and downstream of RT genes were retrieved, clustered at 99.9% sequence identity to remove replicates using CD-HIT^48^ (v4.8.1), and aligned using MAFFT^49^ (v7.505). The resulting alignment was trimmed at the 5′ and 3′ ends to the precise boundaries of the ncRNA as determined by RIP-seq on *Sen*DRT9. These putative ncRNA sequences were clustered at 95% sequence identity using CD-HIT and realigned using mLocARNA^50^ (v1.9.1) with default parameters. The resulting structure-based multiple alignment was used to build and calibrate a covariance model (CM) using the Infernal suite^36^ (v1.1.4). The CMsearch function of Infernal was then used to scan through nucleotide sequences of RT and 1-kb flanking windows, generated by an expanded BLASTp search (max target sequences = 5000) seeded on *Sen*DRT9 and clustered at 85% sequence identity using CD-HIT. The final CM was evaluated for statistically significant co-varying base pairs with R-scape^51^ at an E-value threshold of 0.05, and incorporated a total of 201 homologous DRT9 systems, including *Sen*DRT9.

### ncRNA polyuridine conservation analysis

A window encompassing the polyuridine tracks was extracted from the alignment of DRT9-associated ncRNAs described above, using the *Sen*DRT9 ncRNA as a reference. Next, columns composed of more than 50% gaps were removed with Trimal^52^. A weblogo was then generated from this alignment, before uridine occupancies were calculated for each position with the Biostrings package (v2.74.0) in R and plotted with ggplot2 (v3.5.1).

### *Sen*DRT9 RT-ncRNA complex purification

To purify the His_6_-GST-tagged RT in complex with its ncRNA, *E. coli* BL21-AI cells (NEB) were transformed with the expression vector and cells were grown in LB broth at 37 °C to an OD_600_ of 0.6. Cultures were induced with 0.2 mM IPTG and 0.1% arabinose for overnight expression at 16 °C. Cells were pelleted by centrifugation at 3000 x *g* for 25 min at 4 °C, prior to lysis by sonication in Lysis Buffer I (20 mM Tris-HCl, pH 8.0, 150 mM NaCl, 5 mM MgCl_2_, 5% glycerol, 10 mM imidazole, 0.1% Triton-X-100, 1 mM TCEP). The lysate was clarified by centrifugation at 10,000 x *g* for 30 min at 4 °C. Clarified lysates were passed through a pre-equilibrated Ni-NTA column (Cytiva) using an AKTA purification system (GE Healthcare), then rinsed with 10 column volumes of Wash Buffer (20 mM Tris-HCl, pH 8.0, 150 mM NaCl, 5 mM MgCl_2_, 5% glycerol, 20 mM Imidazole, 1 mM TCEP) prior to elution with Lysis Buffer I supplemented with 300 mM imidazole. The collected fractions were combined and concentrated (100K MWCO PES Spin-X UF concentrator, Corning), prior to size exclusion chromatography using a Sup200 16/600 gel filtration column (Cytiva) pre-equilibrated in Protein Storage Buffer (20 mM Tris-HCl, pH 8.0, 150 mM NaCl, 5 mM MgCl_2_, 5% glycerol, 1 mM TCEP). Fractions containing the complex were pooled and treated with TEV protease overnight before another separation in the same gel filtration column.

Purification of His_6_-2XStrep-SUMO-tagged RT in complex with its ncRNA followed the same culture and induction steps. Supernatants collected after cell lysis were passed through a Strep-Tactin column (IBA) pre-equilibrated in Lysis Buffer II (20 mM Tris-HCl, pH 8.0, 150 mM NaCl, 5 mM MgCl_2_, 5% glycerol, 1 mM TCEP) and eluted with 2.5 mM desthiobiotin (IBA) in Lysis Buffer II. Pooled fractions were concentrated and further purified using a Superose 6 10/300 column (Cytiva). Fractions containing the complex were pooled and treated with Ulp1 protease overnight, before another separation by the same column. The final fractions were pooled, concentrated, aliquoted, flash frozen in liquid nitrogen, and stored at −80 °C.

To purify the dATP homopolymer associated *Sen*DRT9-ncRNA complex, we first incubated 150 nM *Sen*DRT9-ncRNA with 0.9 mM dATP for 30 min at 37 °C. The mixture was analyzed using a Superose 6 10/300 (Cytiva), equilibrated in Protein Storage Buffer (20 mM Tris-HCl, pH 8.0, 150 mM NaCl, 5 mM MgCl_2_, 5% glycerol, 1 mM TCEP).

### Analysis and sequencing of ncRNA co-purifying with RT

Nucleic acids co-purifying with *Sen*DRT9-encoded RT were extracted by treating the RT-ncRNA complex with buffer-saturated phenol (pH 8.5). After vortexing, the sample was centrifuged at 12,000 × *g* for 4 mins, after which the aqueous phase was transferred to a fresh tube and mixed with chloroform. Following an additional round of centrifugation, the recovered aqueous phase was clarified using the Monarch RNA Cleanup Kit (NEB). For analytical RNase/DNase treatments, the purified ncRNA (200 ng) was treated with either 100 Units/mL TurboDNase (Invitrogen) or 250 Units/mL RNase A/T1 (Thermo Scientific) in a 10 μL reaction and incubated for 10 min at 37 °C. Reactions were quenched using 2× RNA Loading Dye (NEB) and heated at 70 °C for 10 min, before electrophoretic separation on a denaturing 10% urea-PAGE gel. Gels were stained with SYBR Gold. RNA-seq library preparation was performed as described above for RIP-seq samples. In brief, RNA was diluted in NEBuffer 2 and heated to 92 °C for 2 min for fragmentation by alkaline hydrolysis, prior to treatment with TURBO DNase (Thermo Fisher Scientific), RppH (NEB), and T4 PNK (NEB). Adapter ligation, reverse transcription, and indexing PCR were performed using the NEBNext Small RNA Library Prep kit. Libraries were sequenced on an Element AVITI in paired-end mode with 150 cycles per end.

### T5 phage gp58 purification

T5 phage-encoded gp58 was purified similarly to the *Sen*DRT9 RT-ncRNA complex, with the following exceptions. Gp58 was expressed as an N-terminal His_6_-2XStrep-SUMO fusion, and was over-expressed in *E. coli* BL21-AI cells (NEB). Cells were induced at OD_600_ of 0.5 with 0.3 mM IPTG and 0.1% Arabinose. After overnight induction at 16 °C, cells were centrifuged at 3000 x *g* for 25 min at 4 °C, sonicated in Lysis Buffer III (20 mM Tris-HCl, pH 8.0, 300 mM NaCl, 5% glycerol), and subjected to affinity chromatography using Strep-Tactin column (IBA). Elution was done in a Lysis Buffer III supplemented with 250 mM desthiobiotin (IBA). The fractions were pooled together, treated with Ulp1 overnight, and passed through a Ni-NTA column equilibrated in the Lysis Buffer III. The flowthrough was collected, concentrated, and was further purified by size exclusion chromatography using a Superdex 75 column (Cytiva) pre-equilibrated in gp58 Storage Buffer (20 mM Tris-HCl, pH 8.0, 300 mM NaCl, 5% glycerol). Fractions were pooled, concentrated, aliquoted, flash frozen in liquid nitrogen, and stored at −80 °C.

### Biochemical poly-dA synthesis assays

Reverse transcription reactions generally contained *Sen*DRT9-encoded RT-ncRNA complexes and nucleotide substrates in Polymerization Buffer I (50 mM Tris-HCl, pH 8.5, 100 mM NaCl, 2 mM MgCl_2_, 5 mM TCEP), at concentrations indicated in associated figure legends. Reactions were initiated by the addition of dATP, typically present at a concentration of 100 µM, and were incubated at 37 °C for 10 min, unless otherwise stated. Radioactive experiments contained trace [α-^32^P]- dATP together with unlabeled (cold) dATP. Reactions were quenched and treated with various nuclease or proteinase reagents prior to electrophoretic separation, as indicated in figure legends. Experiments involving Nuclease P1 digestion were instead incubated in Polymerization Buffer II (50 mM Tris-HCl, pH 7.0, 100 mM NaCl, 2 mM MgCl_2_, 5 mM TCEP) to achieve more optimal conditions for Nuclease P1 cleavage, which loses activity at alkaline pH.

Reactions investigating the fate of unlabeled or radiolabeled [α-^32^P]-dATP substrates were analyzed by denaturing urea-PAGE at 5–10% acrylamide concentrations. Samples were mixed in equal volumes with 2× RNA Loading Dye (NEB) prior to denaturation by heating, followed by electrophoretic separation. Radioactive gels were exposed to a phosphor screen for 5 hours at −20 °C and image using a Typhoon imaging system (GE).

Reactions investigating the fate of the *Sen*DRT9-encoded RT enzyme were analyzed using 10% SDS-PAGE gels. Samples were mixed with 6× SDS Loading Due prior to denaturation by heating and electrophoretic separation. Gels were stained with Coomassie Blue.

### Biochemical competition assays with *Sen*DRT9, ExoI, Nuclease P1, and gp58

Competition assays involving gp58 and ExoI were performed in Polymerization Buffer I. Reactions initially contained 20 nM RT-ncRNA complex and 100 µM dATP, and were first incubated at 37 °C for 10 min, prior to treatment with ExoI in the absence or presence of gp58. ExoI was present at 0.015 Units/µL, and gp58 was titrated into reactions at a range of 0.1–10 µM. Competition assays involving gp58 and Nuclease P1 were performed in Polymerization Buffer II. Reactions initially contained 20 nM RT-ncRNA complex and 100 µM dATP, and were first incubated at 37 °C for 10 min, prior to treatment with Nuclease P1 in the absence or presence of gp58. Nuclease P1 was present at 2 Units/µL, and gp58 was titrated into reactions at a range of 0.1–10 µM.

Reactions were quenched by proteinase K treatment, prior to addition of 2× RNA Loading Dye (NEB), denaturation by heating, and electrophoretic separation by denaturing 5% urea-PAGE.

### Cryo-EM sample preparation

For all samples, Quantifoil copper grids with a mesh size of 300 and R1.2/1.3 hole spacing were glow discharged using a Pelco EsiGlow at 15 mA for 45 seconds. 3 µL of 6 µM purified GST-DRT9 (trimer) or 2.5 µM DRT9 (hexamer) were applied to prepared grids in a Vitrobot Mk IV (ThermoFischer) set to 100% humidity and 4 °C. Grids were subjected to double-sided blotting with a force of 5 for 6s before plunge-freezing in liquid ethane. Clipped grids were loaded into autogrid boxes and stored in liquid nitrogen before imaging.

### Cryo-EM data collection

Both datasets were collected on a Talos Arctica (Thermo Fischer) equipped with a Gatan K3 direct electron detector at Montana State University’s Cryo-EM core facility using SerialEM^53^ (v.4.2.0) under the control of SmartScope^54^ (v0.9.4) for automated data collection. 10,450 and 15,594 micrographs were collected for the hexamer and trimer datasets, respectively. All particles were imaged with a pixel size of 0.9061 Å and apparent magnification of 45,000×. The total dose per exposure was calculated as 59.5 (trimer) and 59.8 (hexamer) electrons/sq.Å/exposure.

### Cryo-EM image analysis, *Sen*DRT9 RT-ncRNA trimer

All image analysis was completed in cryoSPARC^55^ (v4.6.2). After filtering by CTF-fit < 8 Å, and max in-frame motion of 5, 9,814 exposure remained. cryoSPARC Live’s blob picker (50-150 Å particle diameter) was used to identify particles which were then extracted with a box size of 496 px and fourier cropped to 248 px (binx2). 1,878,603 particles were selected from streaming 2D classification and processed to yield a stack of 610,556 particles that were used to generate a 3.7 Å resolution reconstruction without imposing symmetry (**Extended Data Fig. 6b**). This volume was used to generate *de novo* templates for particle picking which yielded 5,235,086 initial picks. 2,530,970 particles were selected from 2-D classification (**Extended Data Fig. 6c**) and fed into a 4 class *ab initio* reconstruction (**Extended Data Fig, 6d**), followed by heterogenous refinement. Non-uniform refinement of the best class produced a 3.7 Å volume containing 2,226,754 particles. After re-extracting unbinned particles and an additional round of 2D classification and 3D sorting by *ab initio* and heterogenous refinement, an initial consensus volume was obtained from 2,107,864 particles. A 5-class 3D classification of these particles without a mask and with force hard classification enabled the removal of additional junk, leaving 1,368,451 particles for further processing. A final round of multi-class *ab initio* reconstruction, followed by heterogeneous refinement (**Extended Data Fig. 6e**) yielded a final stack of 367,640 particles that were refined to 3.02 Å resolution by C3 non-uniform refinement (EMD- 59295) and was used for model building (**Extended Data Fig. 6g–i**).

### Cryo-EM image analysis *Sen*DRT9 RT-ncRNA hexamer

After filtering by CTF-fit < 8 Å resolution, 9,529 exposures remained. 151,584 particles were picked on-the-fly using cryoSPARC Live (100-200 Å particle diameter) and extracted with a box size of 496 px and binned ×4 to expedite processing. A preliminary reconstruction was generated from 48,338 (re-extracted, unbinned) particles that refined to 3.32 Å when D3 symmetry was imposed (3.73 Å, C1). *De novo* templates were produced using this volume and template picking on the full dataset yielded 6,011,113 picks. After two rounds of 2D classification 1,441,369 particles (**Extended Data Fig 7c**) were re-extracted without binning and a box size of 496 px before removing 206,158 duplicate particles. The remaining 1,227,994 particles were subjected to 7-class *ab initio* reconstruction (**Extended Data Fig 7d**) followed by heterogenous refinement. The best class from heterogenous refinement contained 584,077 particles. These particles were re-extracted to obtain better estimates of particle centers before a final round of *ab initio* and heterogenous refinement yielded a stack of 309,844 particles that refined to 2.9 Å resolution by C1 refinement. After volume alignment, imposing D3 symmetry during refinement increased the resolution to 2.59 Å (**Extended Data Fig 7 f-h**). This map (EMD-59293) was used for model building the hexameric DRT9 ribonucleoprotein complex.

### Cryo-EM model building

An initial model for the DRT9 trimer was generated using several copies of an AlphaFold3^37^ structure prediction of the RT monomer and fitting them into the final reconstructions using ChimeraX’s^38^ fit in map command. To model the ncRNA, density maps were submitted to ModelAngelo, along with the ncRNA sequence identified from RIP-seq experiments described above. After building the ncRNA, the complete model for the trimer was refined into the raw map using Phenix RealSpaceRefinement^56^ with reference model and NCS restraints enabled. Iterative editing in COOT^57^ and refinement in Phenix was used to produce the final model (PDB 9NLX). Sharpened maps from Phenix’s anisotropic half-map sharpening tool were used to aid in modelling difficult regions. All outlier residues in the validation report were inspected manually before submission.

The Trimer PDB was used as a starting point for model building of the hexameric form of the complex. After fitting two copies of the trimer structure into the map of the hexamer using ChimeraX’s fit in map command, the model was subjected to RealSpaceRefinement as described above. Iterative editing in COOT and refinement in Phenix were used to produce the final model for the DRT9 hexamer (PDB 9NLV). Sharpened maps from Phenix’s anisotropic half-map sharpening tool were used to aid in modelling difficult regions. All outlier residues were manually inspected before submission.

### Escaper phage isolation

To isolate escaper phages, serial dilutions of phages T5 or Bas37 were mixed in molten soft agar with *E. coli* K-12 strain MG1655 expressing *Sen*DRT9, *Psa*DRT9, or an empty vector control, and plated on LB-agar plates supplemented with chloramphenicol (25 µg/ml). The plates were incubated overnight at 37 °C to allow plaques to form. Individual plaques were picked into 1 mL of MG1655 culture at OD_600_ ∼0.1. Phages forming plaques on *Sen*DRT9 or *Psa*DRT9 plates were inoculated into respective MG1655 cultures expressing *Sen*DRT9 or *Psa*DRT9, while plaques from empty-vector plates were inoculated into MG1655 carrying the empty vector. Cultures were incubated at 37 °C with shaking for 3 hours, and then 800 µL of each culture was transferred to new tubes and mixed with chloroform (5% final volume) to lyse residual bacteria. Cell debris and chloroform were removed by centrifugation for 5 min at 4,000 x *g*, and 5 µL of each phage lysate was propagated further in fresh cultures at OD_600_ ∼0.1. Phages were amplified in this manner for a total of 3 rounds. Escaper phage phenotypes were validated via small-drop plaque assays. T5 escaper phages were isolated from *Sen*DRT9 plates and *Psa*DRT9 plates, and Bas37 escaper phages from *Psa*DRT9 plates; attempts to isolate Bas37 escapers from *Sen*DRT9 were unsuccessful.

### Escaper phage whole-genome sequencing

Phage DNA was isolated by treating 88 µL of phage supernatant with 1 µL of DNase I (200 U/mL, NEB) and 1 µL of RNase A (10 mg/mL, Thermo Fisher Scientific) in 10 µL 1× DNase buffer (NEB). Reactions were incubated at 37 °C for 1 hour, and enzymes were inactivated by heating at 75 °C for 10 min. Phage capsids were digested by adding 1 µL of Proteinase K (20 mg/mL, Thermo Fisher Scientific) and 99 µL of phage lysis buffer (10 mM Tris-HCl, 10 mM EDTA, 0.5% SDS), followed by incubation at 37 °C for 30 min and 55 °C for 30 min. DNA cleanup was performed using Mag-Bind Total Pure NGS beads (Omega), using a bead ratio of 0.9× sample volume.

Phage genomic DNA was tagmented using TnY (a homolog of Tn5) purified in-house following previous methods^58^. 10 ng of purified genomic DNA (gDNA) was tagmented with TnY preloaded with Nextera Read 1 and Read 2 oligos (**Supplementary Table 7**), followed by proteinase K treatment (NEB, final concentration 16 U/mL) and column purification. PCR amplification and Illumina barcoding was done for 13 cycles with KAPA HiFi Hotstart ReadyMix, with an annealing temperature of 63 °C and an extension time of 1 min. The PCR reactions were then pooled and resolved on a gel. A smear from 400– 800 bp was extracted with a Qiaquick Gel Extraction kit (Qiagen) for sequencing on an Element AVITI in paired end mode with 150 cycles per end.

Resulting datasets were processed using cutadapt^39^ (v4.2) to remove Illumina adapter sequences, trim low-quality ends from reads, and filter out reads shorter than 15 bp. Reads were mapped to the T5 (NC_005859.1) genome or Bas37 genome (MZ501089.1), using Bowtie2^59^ (v2.2.1) with default parameters. SAMtools^41^ (v1.17) was used to sort and index alignments, and coverage tracks were visualized in IGV^43^ (v2.17.4). Mutations were identified with breseq^60^ (v.0.39.0) using alignments from Bowtie2 and the same T5 or Bas37 reference genomes as above. T5 and Bas37 escaper mutations are listed in **Supplementary Table 4**.

### Infection response growth curves

*E. coli* K-12 strain MG1655 cells transformed with plasmids encoding WT or MUT *Sen*DRT9 were grown to OD_600_ of 0.2 in LB media supplemented with chloramphenicol (25 µg/mL). 180 µL of each culture was transferred into wells of a 96-well optical plate containing 20 µL of T5 lysate diluted to result in a final MOI of 5 or 0.05, or 20 µL of LB for the uninfected condition. The plate was incubated for 4 hours at 37°C with shaking. OD_600_ values were recorded every 10 min using a Synergy Neo2 microplate reader (Biotek).

### Cell viability assays with trigger and host factor candidates

To assess the effects of *Sen*DRT9 co-expression with phage genes of interest on cell growth, candidate trigger genes identified by escaper screening were cloned onto an expression vector under the control of an arabinose-inducible araBAD promoter. *E. coli* K-12 strain MG1655 cells expressing either WT or MUT *Sen*DRT9 were transformed with trigger plasmids and grown under repressive conditions in LB media supplemented with 2% glucose. Overnight cultures were centrifuged at 4,000 x *g* for 5 min and cell pellets were washed with fresh LB to remove residual glucose. Cells were pelleted again as before and resuspended in fresh LB. 10× serial dilutions of each culture were prepared and 5 µL were spotted on LB agar plates supplemented with chloramphenicol (25 µg/mL), spectinomycin (100 µg/mL), and arabinose (0.2%). Plates were incubated overnight at 37 °C and colonies were counted the next day. Colony forming units (CFU) mL^−1^ were calculated using the following formula: .

To assess cell viability of Δ*sbcB* MG1655 cells expressing *Sen*DRT9, a Δ*sbcB* knock-out strain were transformed with either WT *Sen*DRT9 or MUT *Sen*DRT9. Cells were recovered in 10 mL of LB media at 37 °C for 4 hours. Following recovery, cultures were diluted 1:10 in fresh LB, and 100 µL were plated on LB agar plates supplemented with chloramphenicol and kanamycin. Colony forming units (CFU) mL^−1^ were calculated using the following formula: .

### Nucleotide quantification by LC-MS/MS

*E. coli* K-12 strain MG1655 cells were transformed with empty vector (EV) or *Sen*DRT9 plasmids. Individual colonies were inoculated into LB media supplemented with chloramphenicol (25 µg/mL) and grown overnight at 37 °C with shaking. The following day, cultures were diluted 1:100 in 20 mL fresh LB media and grown to OD_600_ of 0.3. Cultures were split in half and T5 phage was added to one half at an MOI of 2. Uninfected and infected cultures were grown for an additional 45 minutes at 37 °C with shaking. Cells were centrifuged at 4,000 x *g* for 5 min at 4 °C and pellets were washed with 1 mL of cold 1× TBS. Cells were centrifuged again at 15,000 x *g* for 1 min at 4 °C. After supernatants were removed, pellets were flash-frozen in liquid nitrogen and stored at –80 °C.

Cell lysates were prepared as previously described^8^. Flash-frozen pellets were thawed on ice and resuspended in 600 µL of 100 mM sodium phosphate buffer (10:1 mixture of Na_2_HPO_4_ and NaH_2_PO_4_) supplemented with 1 mg mL^−1^ lysozyme. Cells were mechanically lysed using a bead beater (2.5 min × 2 cycles at 4 °C). Lysates were centrifuged at 4,000 x *g* for 15 min at 4 °C and supernatants were transferred to new tubes. Samples were transferred to Amicon Ultra-0.5 3 kDa filter units (Merck Millipore) and centrifuged at 12,000 x *g* for 45 min at 4 °C. Filtrates containing metabolites were collected and stored at –80 °C. Upon retrieval from –80 °C, and prior to use for injection in LC-MS/MS, filtrates were additionally purified by centrifugation at 20,000 x *g* for 20 min at 4 °C. To confirm that each condition contained equal amounts of cellular material, the protein concentration in filtration supernatants was measured by BCA assay (Thermo Fisher Scientific). Protein concentrations varied by < 10% across all samples.

Nucleotide and deoxynucleotide profiling analysis was carried out with ion-paired reversed phase liquid chromatography using an HPLC (1290 Infinity II, Agilent Technologies) coupled to a triple quadrupole mass spectrometer (6495D, Agilent Technologies) with electrospray ionization operated in negative mode. The column was a ZORBAX RRHD Extend-C18 (2.1 × 150 mm, 1.8 µm pore size; 759700-902, Agilent Technologies). 5 µL of the experimental samples were injected. Mobile phase A was 3% methanol (in H_2_O), and mobile phase B was 100% methanol. Both mobile phases contained 10 mM of the ion-pairing agent tributylamine (90780, Sigma Aldrich), 15 mM acetic acid, and 5 µM medronic acid (5191-4506, Agilent Technologies). The LC gradient was: 0-2.5 min 100% A, 2.5-7.5 min ramp to 80% A, 7.5-13 min ramp to 55% A, 13-20 min ramp to 99% B, 20-24 min hold at 99% B. Flow rate was 0.25 mL/min, and the column compartment was heated to 35°C. The column was then backflushed with 100% acetonitrile (0.25 mL/min flow rate 24.05-27 min, followed by 0.8 mL/min flow rate 27.5-31.35 min, and 0.6 mL/min flow rate 31.35-31.50 min) and re-equilibrated with 100% A (0.4 mL/min flow rate 32.25-40 min). The conditions of the MRM transitions for dATP and dTTP were as follows (collision energy (V)): dATP, 490>391.9 (24), 490>158.9 (32); dTTP, 481>383.1 (20), 481>158.8 (36). Fragmentor and CAV were kept constant at 166 V and 4 V, respectively. For other nucleotides and deoxynucleotides measured, conditions of the MRM transitions are shown in **Supplementary Table 8**. In addition, the identity of each compound was confirmed with subsequent injection and acquisition of a pure chemical standard. Data analysis, including peak area integration and signal extraction, was performed with Skyline-daily (v24.1.1.398).

### Nucleoside supplementation assays in plaque assays

Nucleoside supplementation was performed using *E. coli* MG1655 cells transformed with either empty vector or DRT9 expression plasmids. Cultures were grown in LB supplemented with chloramphenicol (25 µg/mL) until OD_600_ reached 0.3. Deoxythymidine, deoxyadenosine, deoxycytidine or deoxyguanosine were added to 4 mL cultures at a final concentration of 20 µM. Cultures were pre-incubated at 37 °C with shaking for 5 min, before adding phage T5 at MOI of 2.5, and grown for another hour. Phages were isolated by adding chloroform to a final concentration of 4%, and samples were centrifuged at 3,000 x *g* for 3 min. Supernatant was used to make 10× serial dilutions used for small-drop plaque assays, performed as described previously. EOP was calculated by spotting the phage on soft agar containing *E. coli* MG1655 cells transformed with an empty vector; the graph shows results from two technical replicates.

### Co-immunoprecipitation Western blot

*E. coli* K-12 strain MG1655 cells were transformed with plasmids encoding either untagged *Sen*DRT9, *Sen*DRT9 with mutated RT catalytic residues (YAAA), N-term 3xFLAG-tagged *Sen*DRT9, or N-term 3xFLAG-tagged *Sen*DRT9 with mutated RT catalytic residues (YAAA), as well as a plasmid encoding C-term V5-tagged gp58. Individual colonies were inoculated into LB media supplemented with chloramphenicol (25 µg/mL), spectinomycin (100 µg/mL) and 2% glucose and grown overnight at 37 °C with shaking. The following day, cultures were diluted 1:100 in 25 mL fresh LB media with continued 2% glucose repression and grown to OD600 of 0.4. Cultures were centrifuged at 4,000 x *g* for 5 min and cell pellets were washed with fresh LB to remove residual glucose. Cells were pelleted again as before, and resuspended in fresh LB with chloramphenicol (25 µg/mL), spectinomycin (100 µg/mL) and 0.2% arabinose, and grown at 37 °C with shaking for 30 min. Cells were centrifuged at 4,000 x *g* for 10 min at 4 °C and pellets were washed with 1 mL of cold 1× TBS. After another round of centrifugation at 15,000 x *g* for 1 min at 4 °C supernatants were removed, and pellets were flash-frozen in liquid nitrogen and stored at –80 °C.

To prepare antibody–bead complexes for immunoprecipitation, Dynabeads Protein G (Thermo Fisher Scientific) were washed 3× in 1 mL IP-MS lysis buffer (50 mM Tris-HCl pH 7.5 at 25 °C, 150 mM NaCl, 5% glycerol, 0.2% Triton X-100). Washed beads were resuspended again in 1 mL IP-MS lysis buffer, combined with anti-FLAG antibody (Sigma-Aldrich, F3165), and rotated for > 3 hours at 4 °C. 75 µL of beads and 10 µL of antibody were prepared per sample. Antibody-bead complexes were washed 2× to remove unbound antibodies and resuspended in IP-MS lysis buffer to a final volume of 75 µL per sample.

Flash-frozen pellets were thawed on ice and resuspended in 1.2 mL IP-MS lysis buffer supplemented with 1× Complete Protease Inhibitor Cocktail (Roche) and 0.1 U/µL SUPERase•In RNase Inhibitor (Thermo Fisher Scientific). Cells were lysed by sonication (2 sec ON, 5 sec OFF, amplitude 20%, 1.5 min total time) and lysates were cleared by centrifugation at 21,000 x *g* for 15 min at 4 °C. Supernatants were transferred to new tubes and protein concentrations were measured by BCA assay (Thermo Fisher Scientific). For each sample, 1 mg of protein was combined with 75 μL of antibody-bead complex and rotated overnight at 4 °C. 50 μL of each cleared lysate was set aside as input control and stored at –80 °C. The next day, each sample was washed 2× with 1 mL cold IP-MS wash buffer 1 (50 mM Tris-HCl pH 7.5 at 25 °C, 150 mM NaCl, 5% glycerol, 0.02% Triton X-100) followed by another 2 washes with cold IP-MS wash buffer 2 (50 mM Tris-HCl pH 7.5 at 25 °C, 150 mM NaCl, 5% glycerol).

For elution, beads were resuspended in 75 μL 2× SDS loading buffer and incubated at 50 °C for 10 min. Input samples were thawed, and both input and IP samples were prepared for SDS-PAGE by combining 15 μL protein with 18.75 μL 2× SDS loading buffer and 3.75 μL 0.1 M DTT. Samples were denatured at 95 °C for 5 min and immediately transferred to ice. Samples were loaded onto 12% Criterion XT Bis-Tris Protein Gels and subjected to electrophoresis in XT MES running buffer at 80 V for 5 min followed by 200 V for 30-40 min until the dye front reached the bottom of the gel. Proteins were transferred to nitrocellulose membranes using iBlot. Membranes were blocked in TBS-T (20 mM Tris-HCl pH 7.5, 150 mM NaCl, 0.1% Tween) containing 5% BSA for 1 hour at room temperature. Membranes were incubated overnight at 4 °C with V5 primary antibody (V5-Tag (D3H8Q), Cell Signaling Technologies) diluted 1:2000 in the blocking buffer. Following three 5-min washes with TBS-T, membranes were incubated with the secondary antibody (mouse anti-rabbit IgG (L27A9)) diluted 1:5000 in a 1:1 mixture of blocking buffer and TBS-T for 1 hour at room temperature. After three 5-min washes with TBS-T, signals were detected using the West Dura Signal kit (Thermo Fisher Scientific) and imaged using chemiluminescence setting on Amersham Imager 600 (GE Healthcare).

### Co-immunoprecipitation mass spectrometry

*E. coli* K-12 strain MG1655 cells were transformed with plasmids encoding either untagged *Sen*DRT9, N-term 3xFLAG-tagged *Sen*DRT9, or N-term 3xFLAG- tagged *Sen*DRT9 with mutated RT catalytic residues (YAAA). Individual colonies were inoculated into LB media supplemented with chloramphenicol (25 µg/mL) and grown overnight at 37 °C with shaking. The following day, cultures were diluted 1:100 in 40 mL fresh LB media and grown to OD_600_ of 0.3. Cultures were split in half and T5 phage was added to one half at an MOI of 5. Uninfected and infected cultures were grown for an additional 45 minutes at 37 °C. Cells were centrifuged at 4,000 x *g* for 10 min at 4 °C and pellets were washed with 1 mL of cold TBS. After centrifuging again at 15,000 x *g* for 1 min at 4 °C and removing supernatants, pellets were flash-frozen in liquid nitrogen and stored at –80 °C.

To prepare antibody–bead complexes for immunoprecipitation, Dynabeads Protein G (Thermo Fisher Scientific) were washed 3× in 1 mL IP-MS lysis buffer (50 mM Tris-HCl pH 7.5 at 25 °C, 150 mM NaCl, 5% glycerol, 0.2% Triton X-100). Washed beads were resuspended again in 1 mL IP-MS lysis buffer, combined with anti-FLAG antibody (Sigma-Aldrich, F3165), and rotated for > 3 hours at 4 °C. 100 µL of beads and 10 µL of antibody were prepared per sample. Antibody-bead complexes were washed 2× to remove unbound antibodies and resuspended in IP-MS lysis buffer to a final volume of 100 µL per sample.

Flash-frozen pellets were thawed on ice and resuspended in 1.2 mL IP-MS lysis buffer supplemented with 1× Complete Protease Inhibitor Cocktail (Roche) and 0.1 U/µL SUPERase•In RNase Inhibitor (Thermo Fisher Scientific). Cells were lysed by sonication and lysates were cleared by centrifugation at 21,000 x *g* for 15 min at 4 °C. Supernatants were transferred to new tubes and protein concentrations were measured by BCA assay (Thermo Fisher Scientific). For each sample, 1 mg of protein was combined with 100 μL of antibody-bead complex and rotated overnight at 4 °C. The next day, each sample was washed 2× with 1 mL cold IP-MS wash buffer 1 (50 mM Tris-HCl pH 7.5 at 25 °C, 150 mM NaCl, 5% glycerol, 0.02% Triton X-100) followed by another 2 washes with cold IP-MS wash buffer 2 (50 mM Tris-HCl pH 7.5 at 25 °C, 150 mM NaCl, 5% glycerol). For elution and on-bead tryptic digestion, beads were resuspended in 80 µL Tris-urea buffer (50 mM Tris-HCl pH 7.5 at 25 °C, 2 M urea, 1 mM DTT) with 0.5 µg/mL sequencing grade modified trypsin (Promega) and incubated 1 hr at 25 °C with regular agitation. Supernatants were transferred to new tubes and beads were resuspended again in 60 µL Tris-urea buffer without trypsin. Supernatants were combined with those from the first elution step. This step was repeated once more, resulting in a total eluate volume of 200 µL per sample. Eluates were centrifuged at 5,000 x *g* for 1 min and supernatants were transferred to new tubes to remove any residual beads. Eluates were flash-frozen in liquid nitrogen and stored at –80 °C prior to further processing.

IP-MS eluates were reduced with 5 mM DTT at 25 °C with 600 rpm agitation for 45 minutes, after which they were alkylated in the dark with 10 mM iodoacetamide (IAA) at 25 °C and 600 rpm for 45 minutes. Samples were then cleaned up using the previously described SP3 protocol^61^. Specifically, 500 µg of an equal mixture of carboxylate-modified hydrophilic beads (Cytiva 45152105050250) and hydrophobic beads (Cytiva 65152105050250) was added to each sample. To induce the binding of proteins to the SP3 beads, 100% ethanol was added to the sample at a 1:1 ratio, and the samples were incubated at 25 °C and 1000 rpm for 15 minutes. After incubation, the beads were washed three times with 80% ethanol and reconstituted in freshly prepared 100 mM ammonium bicarbonate. Samples were digested off the beads using 1 µg of sequencing grade modified trypsin (Promega V5111). After 16 hours of incubation at 25 °C and 600 rpm, samples were taken off the magnetic beads, dried down using a Thermo Savant SpeedVac, and reconstituted in 3% acetonitrile/0.2% formic acid.

Digested peptides from the IP-MS eluates were analyzed on a Waters M-Class UPLC using a 15 cm x 75 µm IonOpticks C18 1.7 µm column coupled to a benchtop Thermo Fisher Scientific Orbitrap Q Exactive HF mass spectrometer. Peptides were separated at a 400 nL min^−1^ flow rate with a 90-minute gradient, including sample loading and column equilibration times. Data were acquired in data-dependent mode using Xcalibur software; each cycle’s 12 most intense peaks were selected for MS2 analysis. MS1 spectra were measured with a resolution of 120,000, an AGC target of 3e6, and a scan range from 300 to 1800 m/z. MS2 spectra were measured with a resolution of 15,000, an AGC target of 1e5, a scan range from 200–2000 m/z, and an isolation window width of 1.6 m/z.

Raw data were searched against a combined reference proteome that included the *E. coli* K-12 strain MG1655 proteome (NCBI RefSeq assembly GCF_000005845.2), T5 phage proteome (NCBI Ref-Seq assembly GCF_000858785.1), and the *Sen*DRT9 RT sequence, using MaxQuant^62^ (v2.0.3.0). One or more unique/razor peptides were required for protein identification. All subsequent analyses were performed in R (v4.4.0). Protein groups with more than five MS/MS spectral counts in at least one condition were retained in the normalized LFQ intensities. A random value between 1 x 10^6^ and 1% of the lowest data point was added as a “pseudocount” to all data points, followed by log_2_ transformation. The data were then used to determine significantly enriched proteins by unpaired two-tailed t-test, with correction for multiple comparisons by the Benjamini-Hochberg method. Significantly enriched interactors were identified using cutoffs of fold-enrichment > 20 and adjusted p-value < 0.05, relative to a control IP from cells expressing untagged WT *Sen*DRT9. IP-MS hits are described in **Supplementary Table 9**.

### Phylogenetic analysis of DRT9 RT and other reverse transcriptases

Sequences of DRT9, a Group II intron RT, KpnDRT2, AbiA, AbiK, AbiP2, and the HIV-1 RT were used to predict monomeric and apo structures of each protein with AlphaFold 3^37^. These predicted structures were then aligned with FoldMason^63^, and the regions corresponding to the Palm and Finger domains were extracted from the alignment. FastTree^33^ was then used to build the phylogenetic tree shown in **Extended Data Fig. 8**.

### Data availability

Next-generation sequencing data will be made available in the National Center for Biotechnology Information (NCBI) Sequence Read Archive at the time of publication. The published genomes used for bioinformatics analyses were obtained from NCBI (**Supplementary Table 1**). Cryo-EM maps and experimentally determined models were deposited to the appropriate public database (i.e. EMDB, PBD) under the accessions EMD-49295, PDB:9NLX (DRT9 Trimer) and EMD-49293, PDB:9NLV (DRT9 Hexamer).

### Code availability

Custom scripts used for bioinformatics, cDIP-seq, RIP-seq, and RNA-seq data analyses are available upon request.

## ACKNOWLEDGMENTS

We thank A.I. Palmieri, S. Kang., and A.J. Robinson for laboratory support; Peter Sims for helpful discussions regarding sequencing data analysis; S. Tavazoie, M. Liu, and P. Oikonomou for generously sharing Keio knockout strains; D. Puleston and R. Sorek for metabolomics support; L.E. Berchowitz for Southern blotting support; the JP Sulzberger Columbia Genome Center for NGS support; and L.F. Landweber for qPCR and gel imager instrument access. Research in the Wiedenheft laboratory is supported by the National Institutes of Health (NIH R35GM134867), the M.J. Murdock Charitable Trust, and the Montana Agricultural Experimental Station. Funding for the Montana State University Cryo-EM Core Facility (RRID:SCR_026324) was contributed by the National Science Foundation (DBI-1828765), the M.J. Murdock Charitable Trust, NIGMS (P30GM140963), and the MSU Office of Research, Economic Development, and Graduate Education. Image processing was performed using the Tempest High Performance Computing System, operated and supported by University Information Technology Research Cyberinfrastructure (RRID:SCR_026229) at Montana State University. S.T. was supported by a Ruth L. Kirch-stein Individual Predoctoral Fellowship (F30AI183830) from the NIH. N.B. was supported by a Ruth L. Kirschstein Individual Predoctoral Fellowship (GM153146) from the NIH and received support from Montana INBRE (P20GM103474). L.M.K. was supported by an NIH Medical Scientist Training Program grant (T32GM145440). M.J. was supported by NIH/NIGMS grant R35GM152258. S.H.S. was supported by NSF Faculty Early Career Development Program (CAREER) Award 2239685, a Pew Biomedical Scholarship, an Irma T. Hirschl Career Scientist Award, a Mallinckrodt Scholarship, the Howard Hughes Medical Institute Investigator Program, and a generous startup package from the Columbia University Irving Medical Center Dean’s Office and the Vagelos Precision Medicine Fund.

## AUTHOR CONTRIBUTIONS

S.T., R.Z., and S.H.S. conceived the project. S.T. and R.Z. performed RIP-seq and cDIP-seq experiments, Southern blotting experiments, and phage genetics experiments. S.T. and M.B. performed metabolomics experiments. S.T. and Y.M. performed proteomics experiments, with supervision from M.J. R.Z. performed nucleoside supplementation and oligo spike-in experiments. N.B. prepared cryo-EM samples, collected and processed cryo-EM data, and built atomic models. S.P. purified SenDRT9-encoded RT-ncRNA complexes and performed all biochemical experiments, with assistance from R.A.W. J.L.R. performed plaque assays with ncRNA mutants, performed phage trigger and host factor experiments, and cloned and tested gp58 variants. L.M.K. cloned and screened DRT9 homologs for activity, isolated escaper phages, and performed and analyzed phage whole-genome sequencing. T.W. performed phylogenetic and bioinformatics analyses and assisted in data interpretation and visualization. D.J.Z. constructed the ncRNA covariance model and performed growth curve experiments. G.D.L. assisted in the design and execution of phage whole-genome sequencing. S.T., R.Z., N.B., S.P., J.L.R., L.M.K., T.W., B.W., and S.H.S. discussed the data and wrote the manuscript, with input from all authors.

## COMPETING INTERESTS

S.H.S. is a co-founder and scientific advisor to Dahlia Biosciences, a scientific advisor to CrisprBits and Prime Medicine, and an equity holder in Dahlia Biosciences and CrisprBits. B.W. is the founder of SurGene LLC and inventor on patent applications related to CRISPR–Cas systems and applications thereof. The remaining authors declare no competing interests.

Correspondence and requests for materials should be addressed to B.W. and S.H.S.

## SUPPLEMENTARY TABLES

Supplementary Table 1 | DRT9-encoded RT homologs in Extended Data Fig. 1a phylogenetic tree.

Supplementary Table 2 | List of DRT9-family immune systems tested in this study.

Supplementary Table 3 | Cryo-EM data collection, refinement and validation statistics.

Supplementary Table 4 | Genotypes of escaper phages that evade DRT9 immunity.

Supplementary Table 5 | Strains used in this study.

Supplementary Table 6 | Description and sequence of plasmids used in this study.

Supplementary Table 7 | Probes and oligonucleotides used in this study.

Supplementary Table 8 | Conditions of MRM transitions from metabolomics measurements.

Supplementary Table 9 | IP-MS hits plotted in Fig. 6d.

