## Extended Data for "Protein-primed DNA homopolymer synthesis by an antiviral reverse transcriptase"

### EXTENDED DATA FIGURES

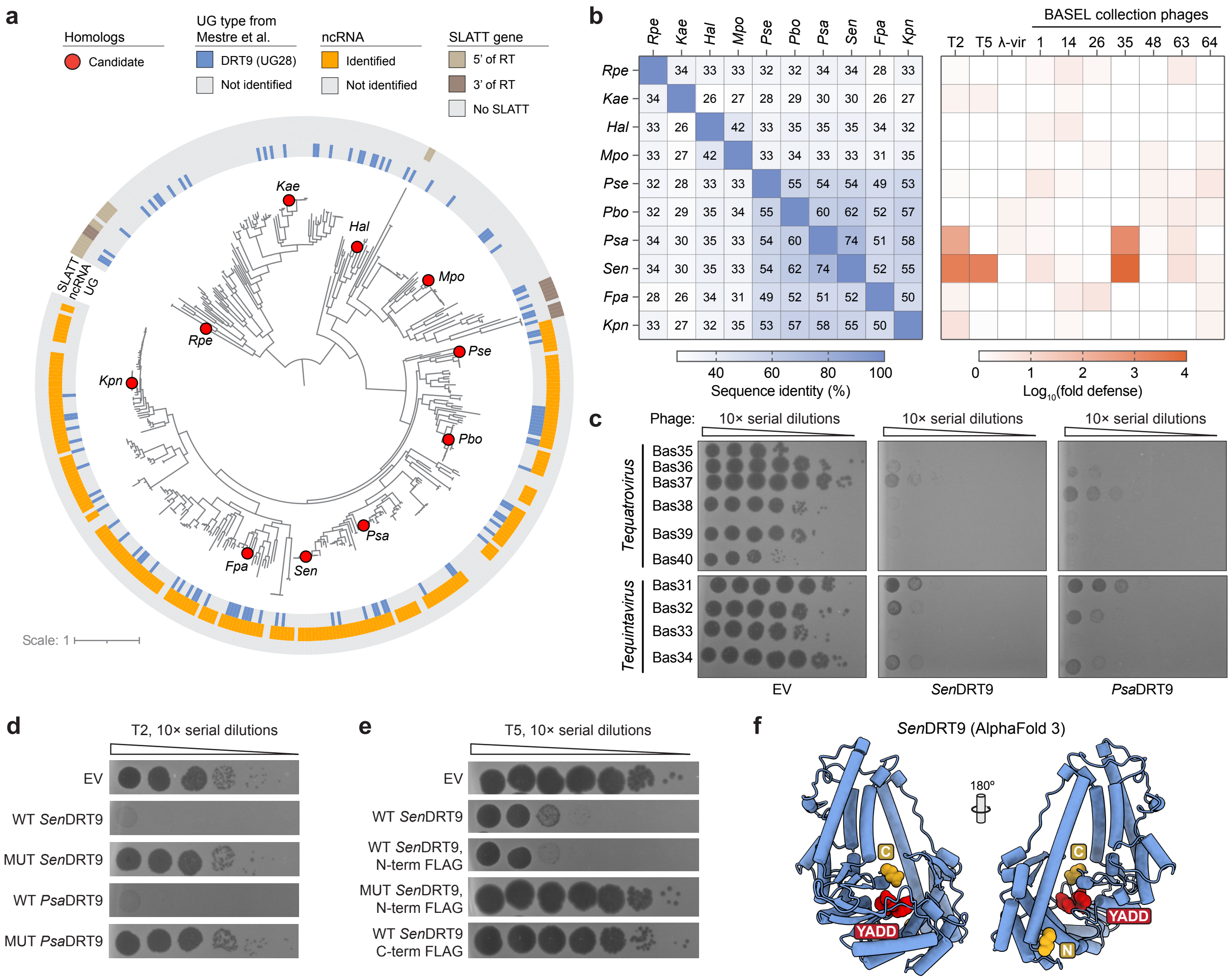


**Extended Data Figure 1 | Screening and identification of active DRT9 immune systems. a,** Phylogenetic tree of DRT9-encoded RT homologs. Outer rings show SLATT protein and ncRNA association within the nearby genomic neighborhood, and the inner ring shows homologs previously identified as falling within the DRT9 (UG28) family[3]. ncRNA association was determined by searching the genomic neighborhood flanking each RT homolog with a covariance model. Homologs selected for experimental testing are indicated with red circles. **b,** Heatmap of pairwise amino acid sequence identity percentages among DRT9-encoded RT homologs tested in this study (left), and heatmap of phage defense activity for the same DRT9 systems tested against 10 diverse *E. coli* phages (right). RT proteins encoded N-terminal FLAG tags for these experiments. **c,** Representative plaque assays demonstrating that *Sen*DRT9 and *Psa*DRT9 exhibit broad defense against phages from the *Tequatrovirus* and *Tequintavirus* genera, as compared to an empty vector (EV) control. **d,** Plaque assays demonstrating that defense activity against T2 phage is completely abolished for both *Sen*DRT9 and *Psa*DRT9 systems encoding catalytically inactive RT mutants (MUT). **e,** Plaque assays demonstrating that *Sen*DRT9-encoded N-terminal FLAG-RT, but not C-terminal RT-FLAG, retains WT defense against T5 phage. **f,** AlphaFold 3 structure prediction of the RT monomer from *Sen*DRT9, highlighting the predicted positions of the N- and C-termini (orange spheres) and YADD active site (red spheres).

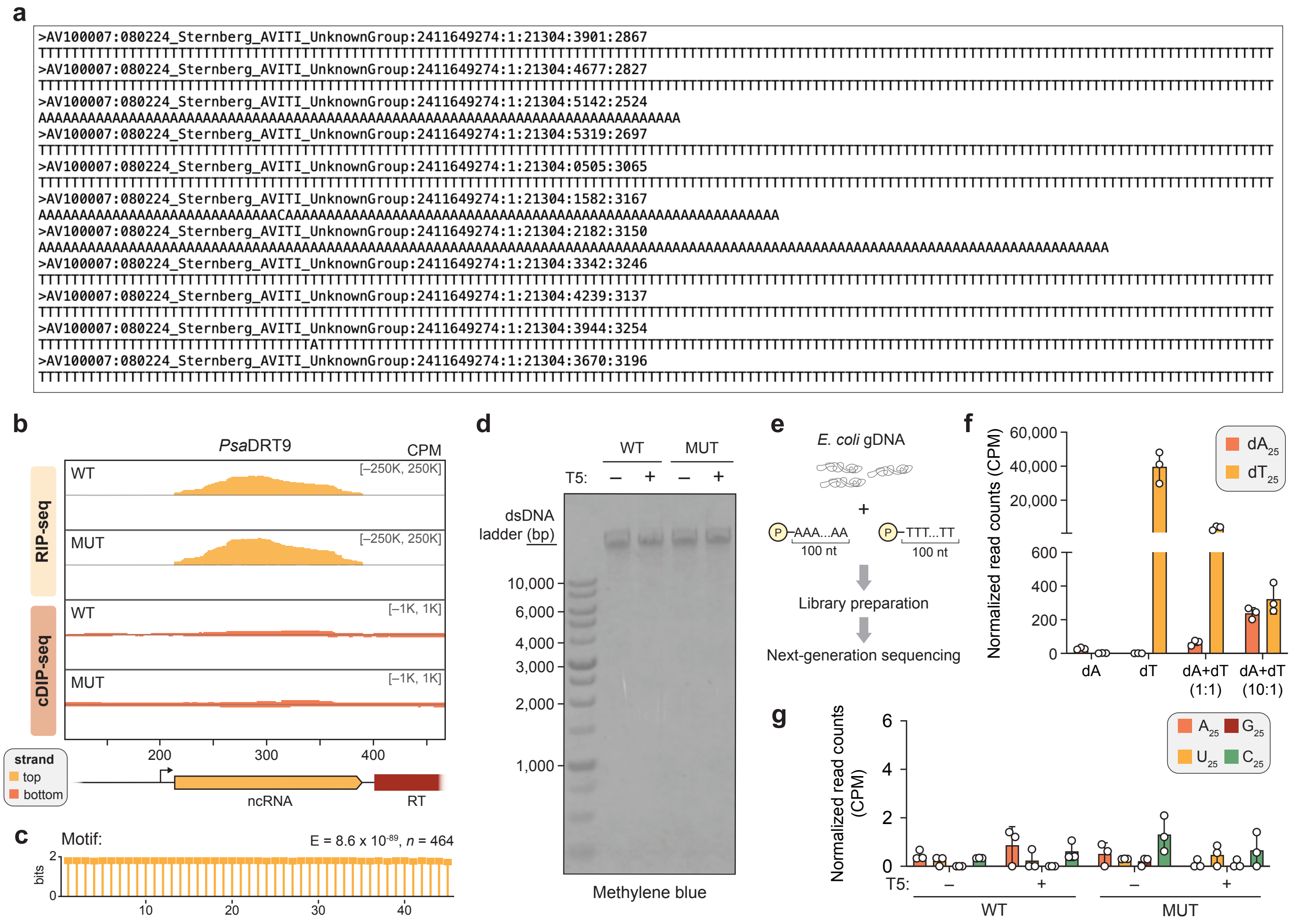


**Extended Data Figure 2 | Discovery and characterization of homopolymeric DNA products elicited by DRT9 immune systems. a,** Raw read calls from an unmapped cDIP-seq dataset of *Sen*DRT9-expressing cells infected with T5 phage, showing uninterrupted strings of poly-dT and poly-dA. **b,** RIP-seq and cDIP-seq coverage tracks (top to bottom) for either WT RT or a catalytically inactive RT mutant (MUT) of *Psa*DRT9, in the presence of T5 phage infection. A schematic of the genomic locus is shown below the graph, and data are normalized for sequencing depth and plotted as counts per million reads (CPM). **c,** MEME analysis results revealing a poly-dT motif enriched in unmapped reads from the WT + T5 cDIP-seq dataset in **b**. *E*, E-value significance of the motif; *n*, number of sites contributing to the motif. **d,** Methylene blue-stained membrane used for the Southern blot shown in **Figure 1h**. **e,** Schematic of oligo spike-in experiment to address potential bias in next-generation sequencing-based detection of poly-dA and poly-dT on the AVITI platform. P, phosphorylated 5′ end. **f,** Bar graph of $dA_{25}$ and $dT_{25}$ counts from total DNA sequencing of oligo spike-in experiments schematized in **e**. The apparent bias against poly-dA capture and sequencing leads to artificially elevated levels of $dT_{25}$-containing reads relative to $dA_{25}$-containing reads. **g,** Bar graph of $dA_{25}$, $dU_{25}$, $dG_{25}$, and $dC_{25}$ counts from total RNA-seq datasets of WT and MUT *Sen*DRT9-expressing cells in the absence or presence of T5 phage infection. These data demonstrate that poly-dA and poly-dT cDNAs are not transcribed, in contrast to the cDNA products of DRT2. Data in **f,g** are shown as mean ± s.d. for n = 3 independent biological replicates.

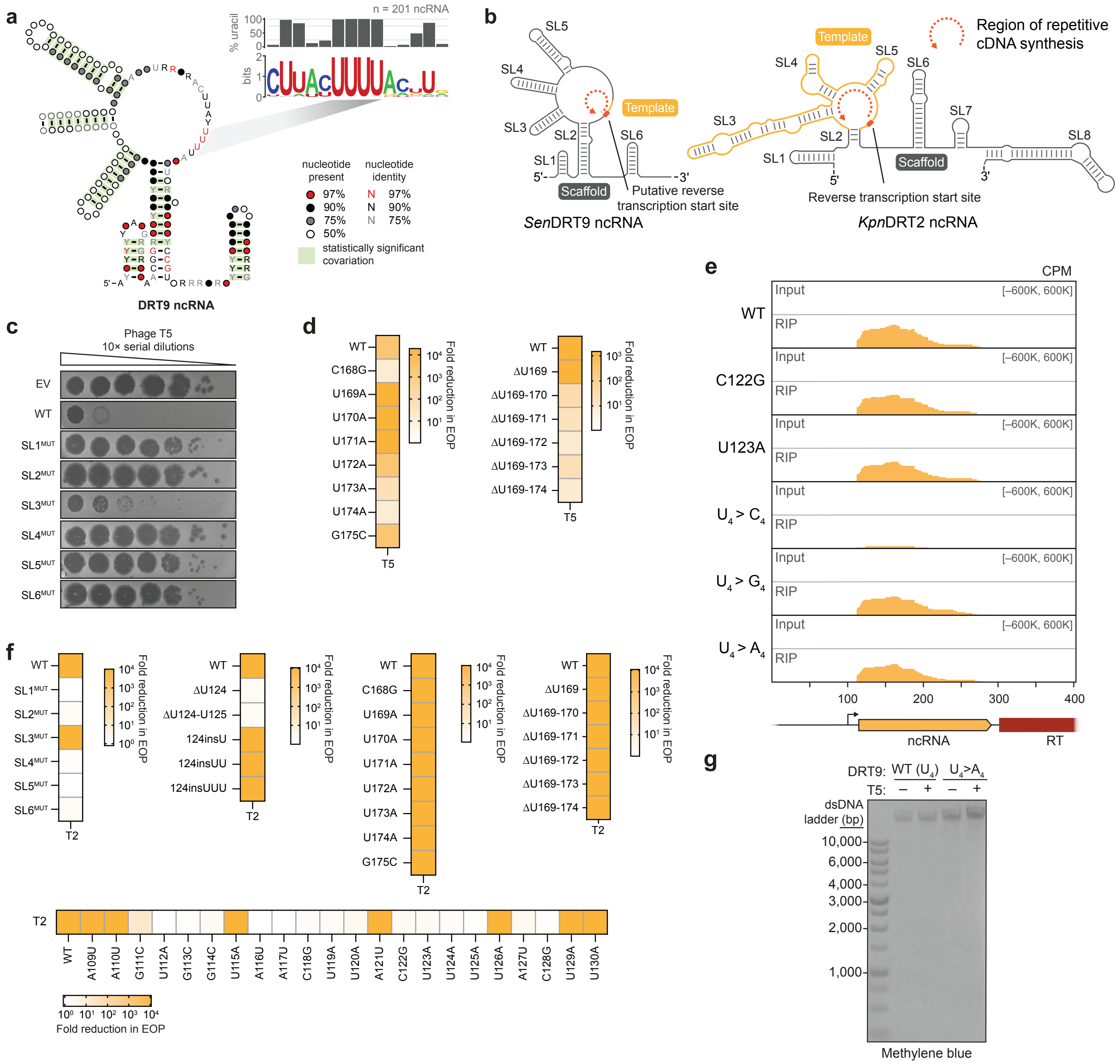


**Extended Data Figure 3 | Additional characterization of ncRNA sequence perturbations on *Sen*DRT9 defense activity.** **a,** Covariance model of the DRT9 ncRNA from an analysis of 201 homologous systems (left), and WebLogo from a multiple sequence alignment centered around the putative U-rich template region (top right). **b,** Comparison of *Sen*DRT9 (left) and *Kpn*DRT2 (right) ncRNAs, highlighting the scaffold (grey) and template (orange) regions. Both ncRNAs template cDNA synthesis from a similar location relative to SL2 (reverse transcription start site, in red), and program repetitive cDNA synthesis across the template region. **c,** Representative plaque assays for the data shown in **Figure 2b**. **d,** Heat map quantifying *Sen*DRT9 defense activity against T5 phage for the indicated ncRNA mutations and deletions within the 3′-proximal SL6 and U-rich region, quantified as the fold reduction in EOP relative to an empty vector (EV) control. Data are shown as the mean of n = 2 technical replicates. **e,** RIP-seq coverage tracks for *Sen*DRT9 with WT ncRNA or the indicated ncRNA mutations in uninfected cells. The bottom three variants are mutated in the putative template region (residues 123–126). A schematic of the genomic locus is shown below the graph, and data are normalized for sequencing depth and plotted as counts per million reads (CPM). **f,** Heat map quantifying *Sen*DRT9 defense activity against T2 phage, for the same ncRNA mutations tested against T5 phage in **Figure 2b–d** and panel **d**. Data are shown as in **d**. **g,** Methylene blue-stained membrane used for the Southern blot shown in **Figure 2f**.

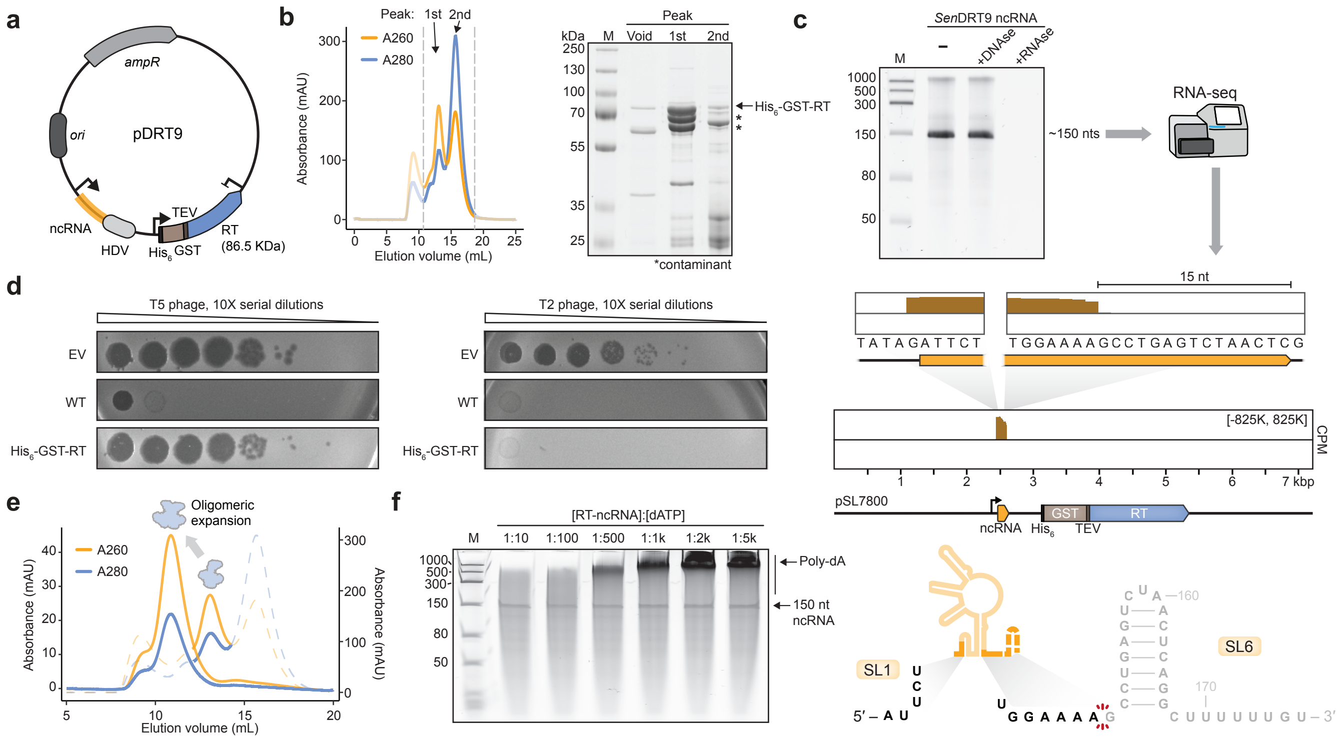

**Extended Data Figure 4 | Purification and characterization of the *Sen*DRT9-encoded RT-ncRNA complex. a,** *E. coli* expression vector design for the *Sen*DRT9-encoded ncRNA and $His_6$-GST-tagged RT. **b,** Size-exclusion chromatogram of the $His_6$-GST-tagged RT-ncRNA complex on a Superdex 200 10/300 column (left), and SDS-PAGE analysis of the void volume and labeled peaks (right). The high $A_{260}$:$A_{280}$ ratio is consistent with a protein-nucleic acid complex. **c,** Denaturing 10% urea-PAGE analysis of the ~150-nt ncRNA species co-purifying with *Sen*RT after RNase or DNase treatment, stained with SYBR Gold (top), and RNA-seq analysis of the ncRNA (middle). The mature ncRNA carries an extraneous 5′-G resulting from the T7 promoter and lacks SL6 at the 3′ end (bottom). This result suggests that SL6 may be involved in transcriptional termination *in vivo* while being dispensable for RT-mediated poly-dA synthesis. **d,** Plaque assay showing loss of *Sen*DRT9 defense activity for a $His_6$-GST-tagged RT variant against T5 phage (left), but not T2 phage (right); EV, empty vector. **e,** Overlaid chromatograms from gel filtration experiments with RT-ncRNA complexes before and after TEV protease treatment of the $His_6$-GST-RT fusion protein, revealing a shift to earlier retention volume and thus increased oligomeric state. The persistent high $A_{260}$:$A_{280}$ ratio suggests that the RT-ncRNA interaction remains intact. **f,** Denaturing 8% urea-PAGE analysis of DNA polymerization assays that contained 150 nM RT-ncRNA complex and increasing ratios of dATP per RT monomer, as indicated. Reactions were incubated at 37 °C for 60 min, followed by proteinase K treatment and phenol-chloroform extraction prior to electrophoretic separation. The gel was stained with SYBR Gold.

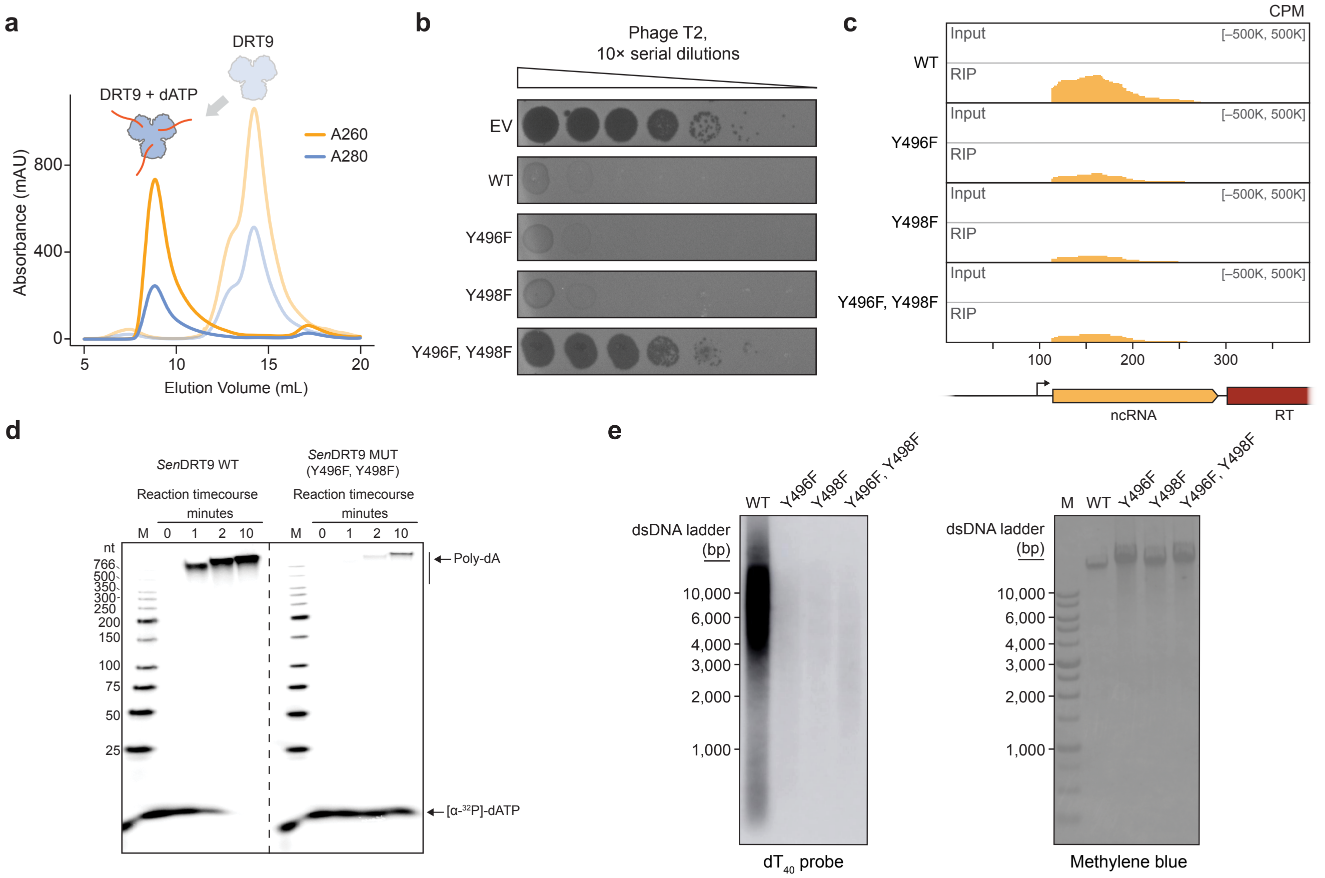


**Extended Data Figure 5 | Roles of C-terminal tyrosine residues in protein-primed reverse transcription and DRT9 phage defense. a,** Overlaid chromatograms from gel filtration analysis of RT-ncRNA complexes with or without dATP incubation, revealing an increased $A_{260}/A_{280}$ ratio and a dramatic shift in retention volume in the presence of dATP. **b,** Plaque assay showing loss of *Sen*DRT9 defense activity against T2 phage for a double Y496F,Y498F mutant, but not single Y496F or Y498F mutants. EV, empty vector. **c,** RIP-seq coverage tracks for *Sen*DRT9 in T5-infected cells with a WT RT compared to single or double Y496F/Y498F mutants, as indicated. A schematic of the genomic locus is shown below the graph, and data are normalized for sequencing depth and plotted as counts per million reads (CPM). **d,** Denaturing 5% urea-PAGE analysis of DNA polymerization assays with a WT RT (left) compared to a double Y496F,Y498F mutant (right). All reactions contained 20 nM RT-ncRNA and 100 µM [α-$^{32}$P]-dATP. M denotes a DNA ladder marker. **e,** Southern blot analysis of total DNA isolated from T5-infected cells expressing *Sen*DRT9 with a WT RT compared to single or double Y496F/Y498F mutants, as indicated. The blot was probed with oligo-$dT_{40}$ (left) to detect poly-dA species. The methylene blue-stained membrane after transfer is shown at right. M denotes a DNA ladder marker.

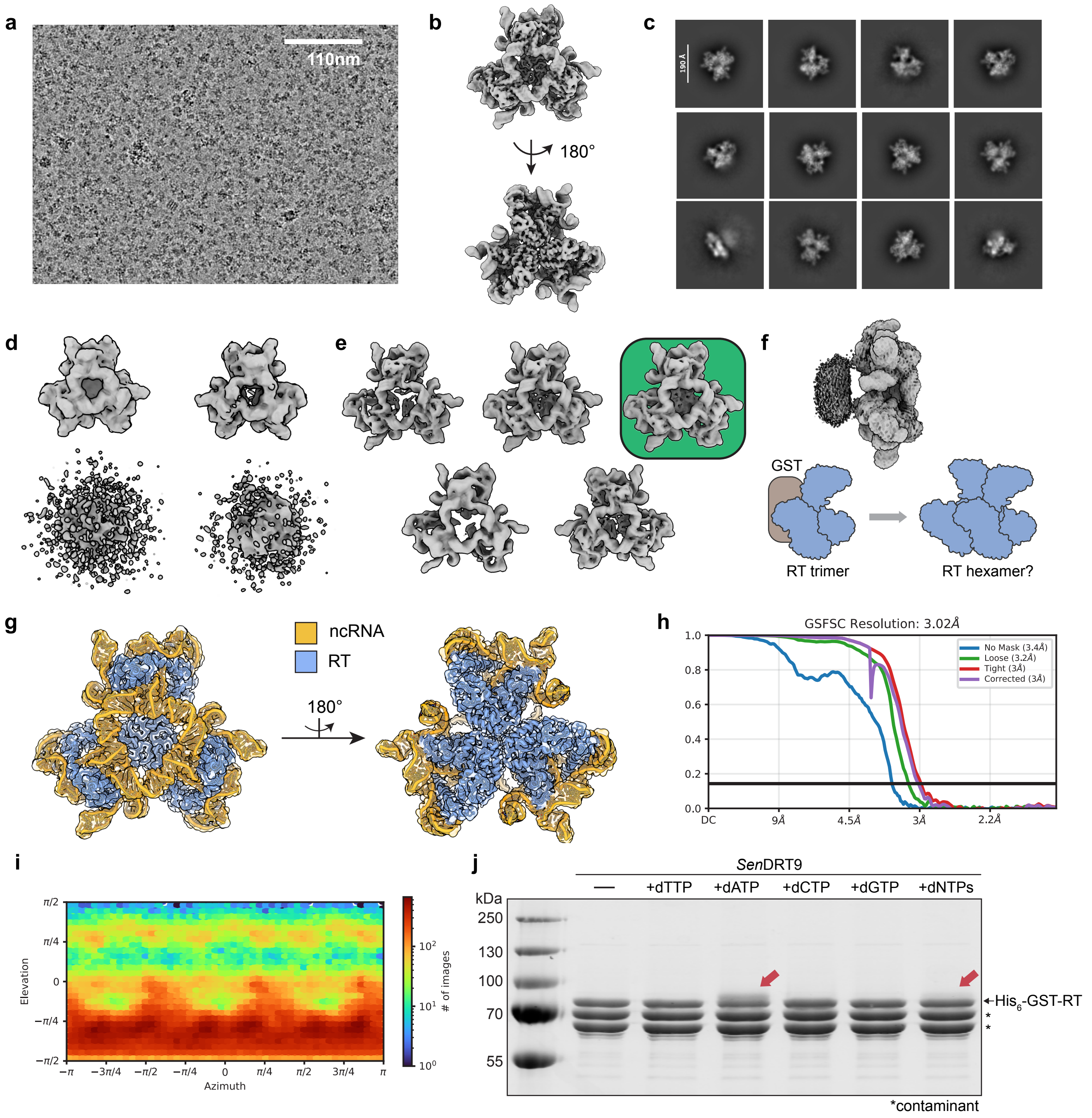


**Extended Data Figure 6 | Cryo-EM analysis and image processing workflow for trimeric *Sen*DRT9 RT-ncRNA complex.** **a,** Representative micrograph from RT-ncRNA complex data collection using the $His_6$-GST fusion protein, revealing a densely-packed field of particles in vitreous ice. **b,** Initial consensus volume after processing blob-picked particles, which refined to 3.7 Å resolution (C3-reconstruction) for *de novo* template generation and subsequent particle picking. **c,** Representative selected 2D classes from 5,235,086 initial particles picked using the template in panel **b**. **d,** Four-class *ab initio* reconstruction from 2,530,970 particles in selected classes to remove junk particles. **e,** Results of a five-class 3D classification without a mask and filtered at 8 Å resolution, with a class similarity of 0.1 to remove remaining poorly aligned particles. The best volume, highlighted by a green background, contained 367,640 particles that were selected for further processing. **f,** View of the final reconstruction shown in **g** shown at high contour (top), revealing noisy density at the N-terminus of each RT monomer corresponding to the $His_6$-GST fusion used to facilitate expression and purification. We hypothesized that the GST fusion could prevent the formation of higher-order oligomers (bottom). **g,** Final (C3) reconstruction that refined to 3 Å resolution and was used for building the trimeric RT-ncRNA complex model. The map is shown with partial transparency and colored according to the model, with RT subunits in blue and ncRNA subunits in orange. **h,** Plot of Cryosparc's gold standard Fourier shell correlation for the map in **g**. **i,** Plot of particle view distribution reveals mild anisotropy but no missing views. **j,** SDS-PAGE analysis of DNA polymerization assays with $His_6$-GST-RT sample, in which RT-ncRNA complexes were incubated with the indicated dNTP(s) for 60 min at 37 °C, before reactions were quenched and resolved electrophoretically. Red arrows indicate preliminary evidence of protein-DNA conjugates in reactions that contained dATP.

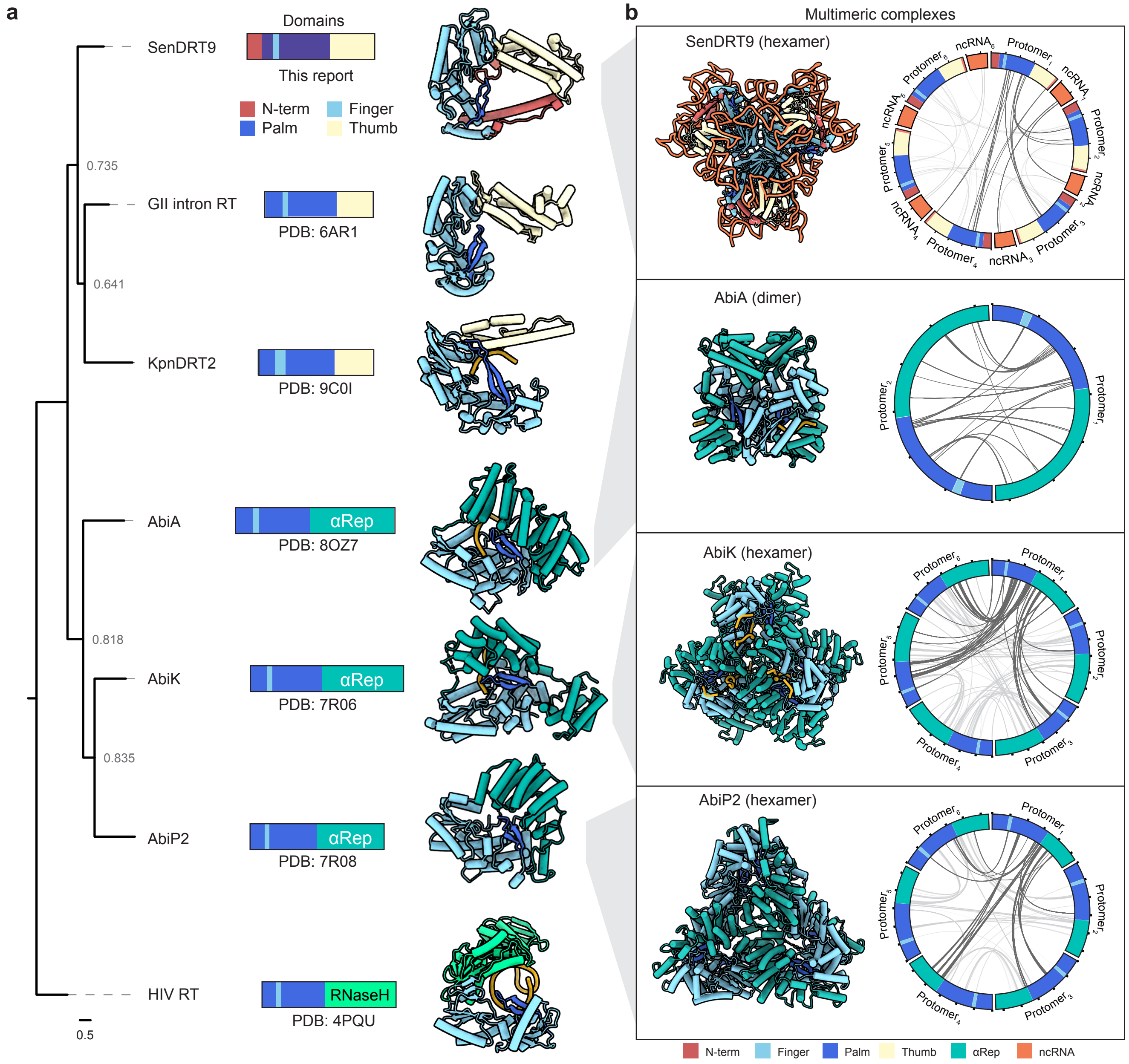


**Extended Data Figure 7 | Comparison of domain composition and 3D structure across evolutionarily diverse RT homologs. a,** Phylogenetic tree of Palm-finger domains of the indicated RT enzymes (left), shown alongside their domain composition (middle) and monomeric structure (right). The tree is rooted to the HIV RT as an outgroup and PDB IDs are shown, alongside the new structure of the *Sen*DRT9-encoded RT presented in this study (top right). **b,** Structural comparison of the indicated bacterial RT multimeric complexes (left), with the hexameric *Sen*DRT9 RT-ncRNA complex shown at the top. AbiA, AbiK, and AbiP2 rely exclusively on protein-protein interactions, whereas the ncRNA plays a crucial role in DRT9 hexamer assembly, as highlighted in the interactome plots (right). Protein domains and ncRNA are colored according to the legend shown at the bottom.

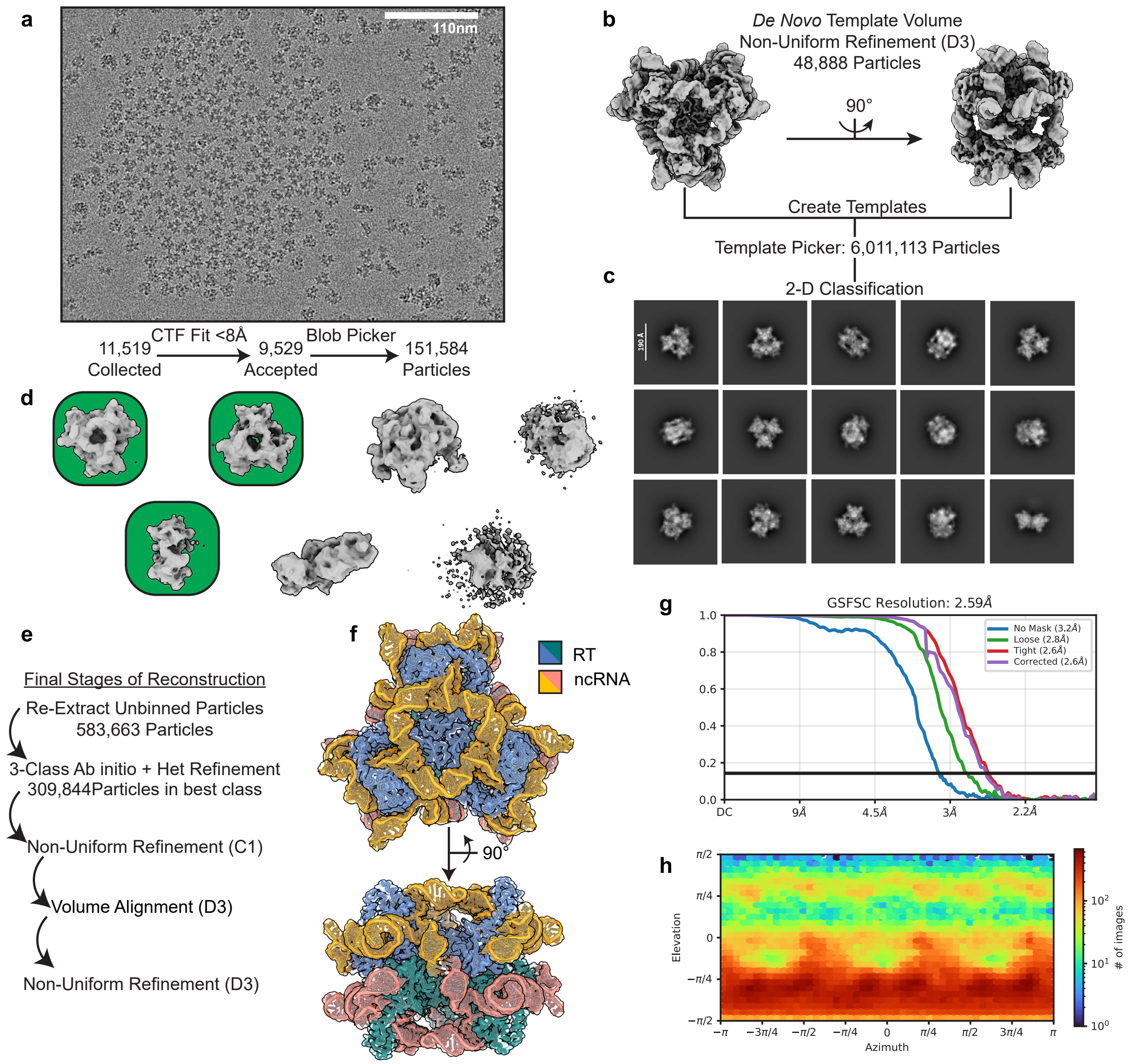


**Extended Data Figure 8 | Cryo-EM analysis and image processing workflow for hexameric *Sen*DRT9 RT-ncRNA complex. a,** Representative micrograph from the RT-ncRNA hexameric complex, revealing a monolayer of evenly distributed particles. **b,** Initial reconstruction from on-the-fly image analysis that was used to generate *de novo* templates used for particle picking. **c,** Representative 2D Classes from 6,011,113 template-picked particles. **d,** Initial volumes from a 7-class *ab initio* reconstruction of 1,411,369 particles. Selected volumes for downstream processing are shown with a green background. **e,** Final steps used to sort selected particles in **d** to yield the final reconstruction. **f,** Final (D3) reconstruction of the *Sen*DRT90-encoded RT-ncRNA hexamer that refined to 2.6 Å and was used for model building. The map is shown as a partially transparent surface, with RT subunits shown in blue/cyan and ncRNA subunits shown in orange/salmon. **g,** Plot of Cryosparc's gold standard Fourier shell correlation for the map in **f**. **h,** Plot of particle view distribution reveals mild anisotropy but no missing views.

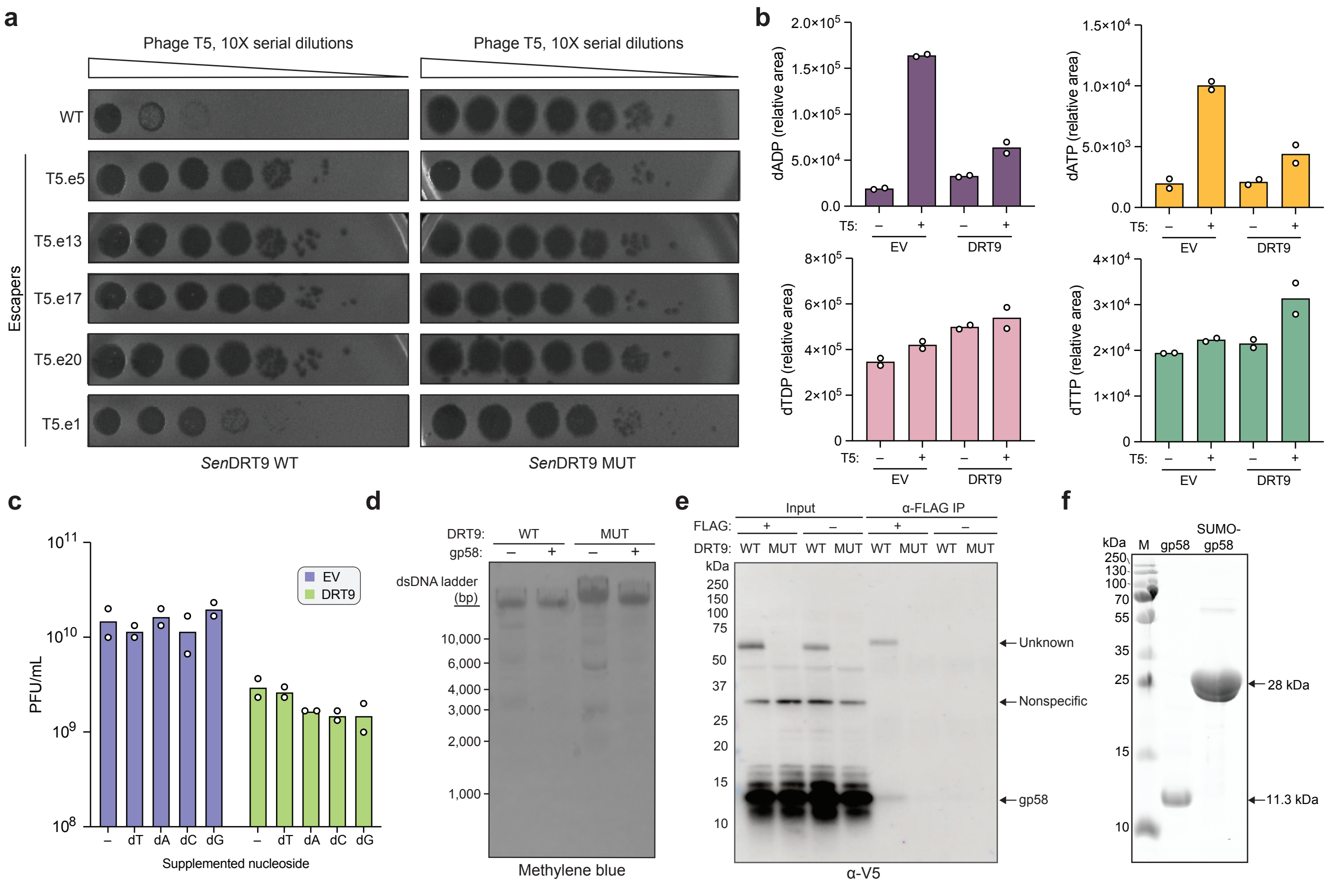


**Extended Data Figure 9 | Activation of DRT9 immunity by phage-encoded factors**. **a,** Representative plaque assays with WT or escaper variants of T5 phage, tested in strains expressing either the WT or RT-inactive (MUT) *Sen*DRT9 system. **b,** Bar graphs of relative nucleotide levels for the indicated species, quantified by LC-MS/MS, in lysates from cells expressing *Sen*DRT9 or an empty vector (EV) control in the absence or presence of T5 phage infection. Data are shown as the mean for n = 2 independent biological replicates. **c,** Bar graphs of plaque forming units per mL (PFU/mL) of T5 phage lysates after liquid culture infection of cells expressing *Sen*DRT9 or an empty vector (EV) control, in the presence of the indicated supplemented nucleosides. Data are shown as the mean for n = 2 independent biological replicates. **d,** Methylene blue-stained membrane of total DNA from cells expressing WT or MUT *Sen*DRT9 with (+) or without (–) gp58 induction, for the Southern blot shown in **Figure 5d**. **e,** Western blot analysis of *Sen*RT co-immunoprecipitation with gp58. Immunoprecipitation was performed with a FLAG antibody on lysates from cells co-expressing *Sen*RT (WT or MUT, FLAG-tagged or untagged) with V5-tagged gp58. Together with non-immunoprecipitated controls (Input), samples were analyzed by Western blot using a V5 antibody. The unique presence of a band corresponding to gp58 in the IP eluate from cells expressing WT *Sen*DRT9 indicates a DNA-dependent interaction between the RT and gp58. **f,** SDS-PAGE analysis of gp58 purification, showing protein purity before and after protease-based removal of the N-terminal $His_6$-SUMO tag. M denotes a protein ladder marker.
