## Supplementary Table 3 for "Protein-primed DNA homopolymer synthesis by an antiviral reverse transcriptase"

**Supplementary Table 3: Cryo-EM data collection, refinement and validation statistics**

|  | #1 GST-DRT9 Trimer  (EMDB-49525)  (PDB 9NLX) | #2 DRT9 Hexamer, Pre-Polymerization State  (EMDB-49523)  (PDB 9NLV) |
| --- | --- | --- |
| **Data collection and processing** |  |  |
| Magnification | 45,000 | 45,000 |
| Voltage (kV) | 200 | 200 |
| Electron exposure (e–/Å^2^) | 59.5 | 59.8 |
| Defocus range (μm) | -0.8, -2.5 | -0.8, -2.5 |
| Pixel size (Å) | .9061 | .9061 |
| Symmetry imposed | C3 | D3 |
| Initial particle images (no.) | 5,235,086 | 6,029,711 |
| Final particle images (no.) | 367,640 | 309,844 |
| Map resolution (Å)  FSC threshold | 3.02  0.143 | 2.59  0.143 |
| Map resolution range (Å) | 2.6-6.0Å | 2.0-4.5 |
| **Refinement** |  |  |
| Initial model used (PDB code) | AlphaFold3 | AlphaFold3 |
| Map sharpening methods  Refinement Package  Refinement Method  **Model composition**  Non-hydrogen atoms  Protein residues  Nucleotide Residues  Ligands | Phenix Half-Map  Phenix  RealSpaceRefinement  19,647  1,458  411  0 | Phenix Half-Map  Phenix  RealSpaceRefinement  40,284  2,294  828  0 |
| ***B* factors (Å^2^)**  Protein  Nucleotide  Ligand | 31.08  4.85  N/A | 33.62  15.93  N/A |
| **R.m.s. deviations**  Bond lengths (Å)  Bond angles (°) | 0.002  0.627 | 0.002  0.573 |
| **Validation**  MolProbity score  Clashscore  Poor rotamers (%) | 1.54  3  1 | 1.54  5.52  1 |
| **Ramachandran plot**  Favored (%)  Allowed (%)  Disallowed (%) | 98  2  0 | 98  2  0 |
